# Vesicles driven by dynein and kinesin exhibit directional reversals without external regulators

**DOI:** 10.1101/2022.09.27.509758

**Authors:** Ashwin I. D’Souza, Rahul Grover, Gina A. Monzon, Ludger Santen, Stefan Diez

**Author notes:** equal contribution (Ashwin I. D’Souza, Rahul Grover, Gina A. Monzon).

## Abstract

Intracellular transport along cytoskeletal filaments propelled by molecular motors ensures the targeted delivery of cargoes to their destinations. Such transport is rarely unidirectional but rather bidirectional, including intermittent pauses and directional reversals owing to the simultaneous presence of opposite-polarity motors. It has been unclear whether such a complex motility pattern results from the sole mechanical interplay between opposite-polarity motors or requires external regulators. Here, we addressed this outstanding question by reconstituting cargo motility along microtubules *in vitro* by attaching purified Dynein-Dynactin-BICD2 (DDB) and kinesin-3 (KIF16B) to large unilamellar vesicles. Strikingly, we found that this minimal system is sufficient to recapitulate runs, pauses and reversals similar to *in vivo* cargo motility. In our experiments, reversals were always preceded by vesicle pausing and the transport directionality could be tuned by the relative numbers of opposite-polarity motors on the vesicles. Unexpectedly, during all runs the vesicle velocity was not influenced by the presence of the opposing motors. To gain mechanistic insight into bidirectional transport, we developed a mathematical model which predicts that low numbers of engaged motors are critical to transition between runs and pauses. Taken together, our results suggest that motors diffusively anchored to membranous cargo transiently engage in a tug-of-war during pauses where stochastic motor attachment and detachment events can lead to directional reversals without the necessity of external regulators.

## Introduction

Intracellular organelles such as endosomes, synaptic vesicles and lipid droplets are transported as cargoes along polarized microtubule filaments by minus-end directed cytoplasmic dynein (referred to as ‘dynein’) and plus-end directed kinesin motors. Multiple copies of dynein and kinesin are simultaneously present on individual cargoes leading to bidirectional motion ^1–10^. This motion is characterized by fast runs in either direction and frequent directional reversals. Despite these reversals, the direction of cargo transport is biased towards the intended intracellular destination. The origin of directional reversals and their regulation to achieve targeted transport remain poorly understood.

The transport direction of cargoes undergoing reversals can be potentially biased by a variety of factors. The primary mode of action of these biasing factors would be to differentially support or impede the teams of dyneins and kinesins associated with the cargo. As such, cargo adaptors that either activate dynein (e.g. BICD2^11,12^, ninein^13^) or kinesin (e.g. nesprin-4 activation of kinesin-1^14^) have been identified as biasing factors. Moreover, certain microtubule-associated proteins (MAPs) have been shown to exhibit differential influences on dynein and kinesin ^1,15–18^. In view of the numerous factors that can influence the behavior of motor-cargo systems, highly controlled, cell-free *in vitro* reconstitution assays are essential to probe the interplay between dynein and kinesin on cargoes and investigate the effect of candidate biasing factors.

Previously, a number of *in vitro* reconstitution experiments with artificial assemblies of dynein and kinesin linked to DNA origami chassis, short stretches of double-stranded DNA or glass surfaces have been reported ^19–22, 23^. However, all of them were limited in their capability to recapitulate key features of intracellular motility, such as fast unidirectional transport and directional reversals. The assemblies either remained stationary or moved at markedly low velocities suggesting a constant tug-of-war between dynein and kinesin. *In vivo*, tugs-of-war are indeed hypothesized to occur ^1,3,6–9^ and spatial elongations, for example of endosomes isolated from *Dictyostelium discoideum*, along the microtubules have been regarded a signature of the underlying opposing forces ^9^. However, these tugs-of-war are intermittent and last only for short periods (one to two seconds) before fast unidirectional transport resumes. The lack of fast transport and directional reversals with artificial assemblies *in vitro* indicated that recapitulating features of intracellular cargo motility required an additional activity that, in turn, reciprocally regulated the activities of dynein and kinesin ^19^. Such regulation would prevent simultaneous force generation by the opposite polarity motors, thereby reducing the prospect of a constant tug-of-war. Nevertheless, one key difference between previous artificial assemblies and intracellular cargoes is the nature of the cargo itself. The artificial assemblies rigidly coupled fixed motor compositions to each other. Intracellular cargoes, on the other hand, are usually vesicular structures made of lipid membranes that allow for motor diffusion. Properties of lipid membranes, such as geometry and fluidity, can influence transport characteristics. For example, spherical vesicles driven by multiple myosin Va motors move faster than single myosin Va motors ^24,25^ and the transport efficiency of multiple lipid-anchored kinesin-1 motors is reduced in fluid membranes^26^. So far, it has been unclear whether replacing artificial assemblies with vesicular cargoes would recapitulate the bidirectional features of intracellular cargo motility.

Here, we develop a well-defined reconstitution assay with purified dynein and kinesin-3 as motors and small unilamellar vesicles of defined phospholipid composition as cargo. We show that vesicles driven by dynein and kinesin-3 exhibit the features of intracellular cargo motility, namely fast minus- and plus-end directed runs, intermittent pauses as well as directional reversals. We observe that the simultaneous presence of dynein and kinesin-3 do not affect the velocity of the vesicles during unidirectional runs but increase the frequency and duration of pauses. We find that directional reversals are often preceded by a tug-of-war which manifests as deformations of larger vesicles. In agreement with numerical simulations, our results suggest that motors diffusively anchored on vesicles do not hinder each other significantly during runs, but engage in a tug-of-war during intermittent pauses where stochastic fluctuations in the number of engaged motors can lead to directional reversals without the necessity of external regulators.

## Results

### Purified Dynein-Dynactin-BICD2 complexes and KIF16B motors are processive

We assembled a toolkit comprising functional minus- and plus-end directed motors. For the minus-end directed motors, we purified native *H. sapiens* dynein and dynactin complexes from Human Embryonic Kidney 293 (HEK293) cells as well as *M. musculus* bicaudal 2 (BICD2) truncated to first 594 amino acids (BICD2N594, BICD2N594-eGFP) from *E. coli*, as dynein activator (**Fig. 1a**).The dynein-dyactin-BICD2N594-eGFP (DDB-eGFP; 1:2:1.5) complex was active and moved along surface-immobilized microtubules (**Fig. 1b** left panel, **Supplementary video 1**) with a median instantaneous (frame-to-frame) velocity of -1.46 ± 1.63 µm/s (± IQR, **Fig. 1c** upper panel) and a median run length of 3.29 ± 4.43 µm (± IQR, **Extended Data Fig. 1a**). For the plus-end motor, we purified full-length *H. sapiens* KIF16B (with and without a C-terminal eGFP tag, **Fig. 1a**) from *Spodoptera frugiperda* (Sf9+) cells. KIF16B is a kinesin-3 motor responsible for the anterograde motility of early endosomes ^27,28^. It contains a Phox homology (PX) domain at its C-terminus, which can directly bind to membranes containing phosphatidylinositol-3-phosphate (PI3P) phospholipids ^28,29^. KIF16B-eGFP motors exhibited processive motility along surface-immobilized microtubules (**Fig. 1b** right panel, **Supplementary video 2**) with a median velocity of 0.80 ± 0.63 µm/s (± IQR, **Fig. 1c** lower panel) and a median run length of 0.63 ± 0.68 µm (± IQR, **Extended Data Fig. 1b**). KIF16B, along with other members of the kinesin-3 family, is hypothesized to largely exist as autoinhibited monomers in cells, and undergoes dimerization only when recruited to cargoes ^30^. However, our recombinantly expressed KIF16B was active and showed processive motility indicating that at least a subset of motors can dimerize without a cargo.

**Figure 1.**
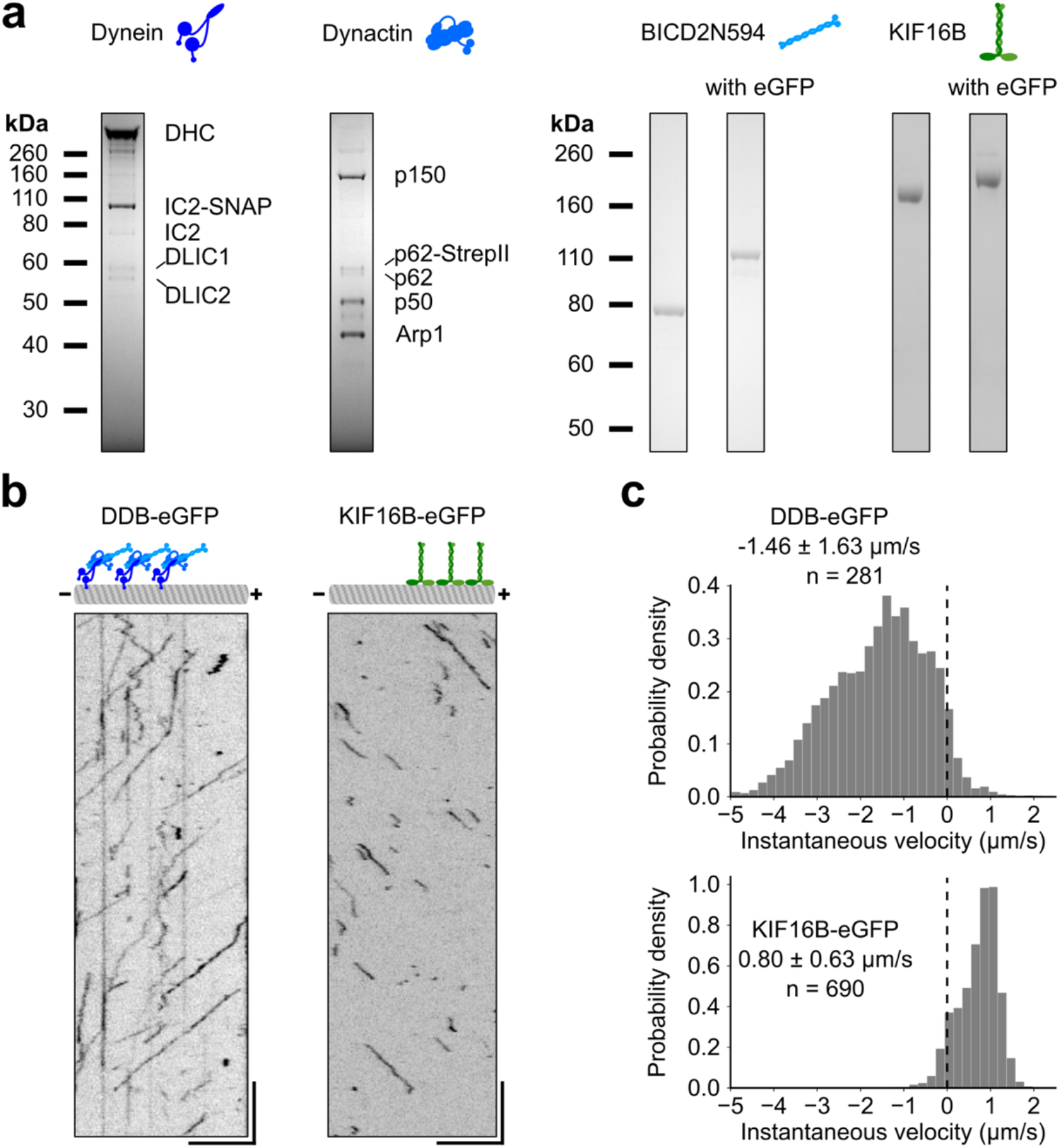
Purified Dynein-Dynactin-BICD2N594 (DDB) complexes and KIF16B motors are processive. **a)** SDS–polyacrylamide gel electrophoresis (SDS-PAGE) of purified Dynein, Dynactin and BICD2N (truncated to first 594 aa., untagged and tagged with an enhanced green fluorescent protein (eGFP)) and Kinesin-3 (KIF16B, untagged and tagged with eGFP). Identity of DHC, IC2, p150, p62, p50 and Arp1 were confirmed with Western blotting using appropriate antibodies (see Materials and Methods). **b)** Representative kymograph of single DDB complexes visualized with BICD2N-594-eGFP (DDB-eGFP, left panel) and single KIF16B-eGFP molecules (right panel). Motility was observed under a TIRF microscope at 28 °C in the presence of 2.5 mM MgATP. Position-time data was obtained by tracking single complexes/molecules, correcting for drift with 0.1 µm fluorescent beads and calculating the position along the microtubule. Scale bars: vertical 5 s, horizontal 5 µm. **c)** Histograms of instantaneous velocity (velocity between consecutive frames) of single DDB complexes (upper panel) and single KIF16B-eGFP motors (lower panel). Data is reported as median ± inter-quartile range (IQR). n represents the number of single-molecules/complexes. The probability density is defined as the number of counts per bin divided by the product of total count and bin width. The integral over the histogram is equal to one.

### DDB-KIF16B-vesicles exhibit directional reversals *in vitro*

To observe the motility of vesicles driven by either or both opposite-polarity motors along microtubules, we first prepared large unilamellar vesicles (diameter of 132.5 ± 48.5 nm) containing DGS-NTA(Ni) and phosphoinositol-3-phoshpate (PI3P) to attach DDB (dynein-dyactin-BICD2N594-8xHis) and KIF16B, respectively (Materials and Methods, **Extended Data Fig. 2a**, and **Extended Data Table 1**). We recorded the motility of motor-bound vesicles, diluted in imaging buffer, along polarity-marked microtubules^31^ under a Total Internal Reflection Fluorescence (TIRF) microscope (**Fig. 2a**). We first characterized the unidirectional vesicle motion towards the minus-end with DDB. DDB-vesicles were prepared by incubating vesicles with saturating amounts (7-fold higher than DGS-NTA(Ni) concentration) of BICD2N594-8xHis followed by the addition of pre-incubated dynein-dynactin (1:2) complexes (38 nM DDB). DDB-vesicles moved over long (> 10 µm) distances towards the minus-end (**Fig. 2b** upper panel, **Supplementary video 3**). At this concentration of motors, DDB-vesicles often traverse the entire length of the microtubule. However, vesicle motility was frequently interrupted by pauses leading to a peak around zero in the histogram of instantaneous velocities and yielded a median velocity of -0.40 ± 0.94 µm/s (± IQR, **Fig. 2b** lower panel).

**Figure 2.**
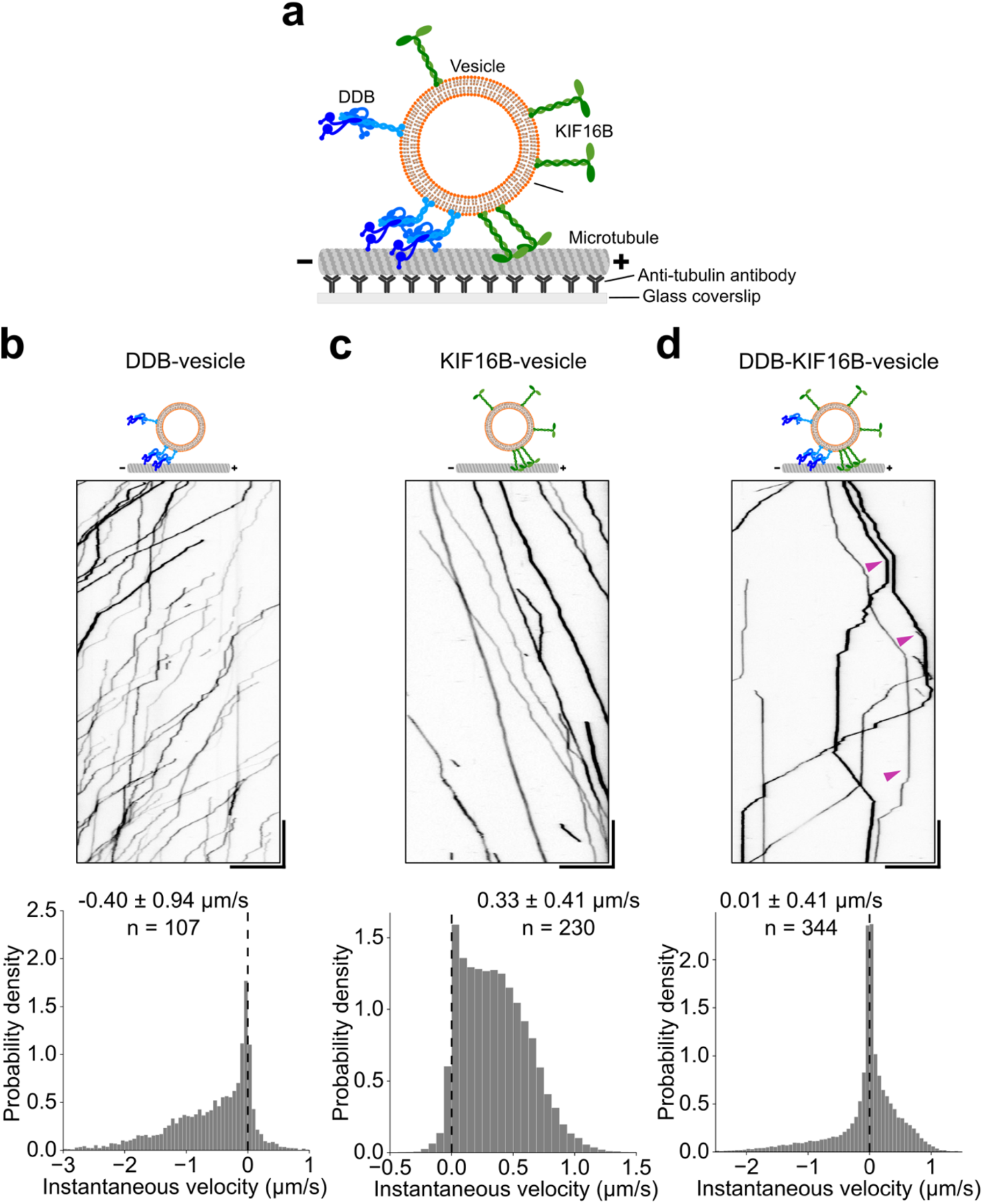
DDB-KIF16B-vesicles exhibit directional reversals in vitro. **a)** Schematic diagram of vesicle motility assay. **b-d)** Kymographs (upper panels) and velocity histograms (lower panels) of DDB-vesicle motility (b), KIF16B-vesicle motility (c), and DDB-KIF16B-vesicle motility (d) on polarity-marked microtubules. Magenta arrowheads mark vesicles that exhibit directional reversals. Motility was observed at 28 °C in the presence of 2.5 mM MgATP. Position-time data was obtained by tracking vesicles, correcting for drift with 0.1 µm fluorescent beads, and calculating the position along the microtubule. Scale bars: vertical 10 s, horizontal 10 µm. Numerical values are reported as median ± inter-quartile range (IQR). n represents the number of vesicles. Data from two independent experiments was combined to create the histograms.

We then characterized the unidirectional vesicle motion towards the plus-end with KIF16B. KIF16B-vesicles were prepared by incubating vesicles with 25 nM KIF16B. KIF16B-vesicles also exhibited robust motility over long distances (> 10 µm; **Fig. 2c** upper panel, **Supplementary video 4**) often reaching the end of the microtubule. Processive runs of KIF16B-vesicles were also interrupted by pauses leading to a peak around zero in the histogram of instantaneous velocities and yielded a median velocity of 0.33 ± 0.41 µm/s (± IQR, **Fig. 2c** lower panel).

Next, we tested if vesicles would still move unidirectionally even in the presence of both DDB and KIF16B (dual-motor vesicle assay). DDB-KIF16B-vesicles were prepared by first incubating vesicles with BICD2N594 (7-fold excess with regard to DGS-NTA(Ni)) and 25 nM KIF16B, followed by the addition of dynein-dynactin (1:2) complexes (38 nM). We observed unidirectional vesicles either moving towards the minus-end or the plus-end as well as vesicles exhibiting directional reversals from minus-end directed motion to plus-end directed motion or vice-versa (**Fig. 2d** upper panel, magenta arrowheads, **Supplementary video 5**). As in the case of DDB-vesicles and KIF16B-vesicles, directed runs of DDB-KIF16B-vesicles were often interrupted by pauses. In fact, this pausing behavior appeared to be enhanced in case of DDB-KIF16B-vesicles as indicated by a major peak around zero in the histogram of instantaneous velocities and yielded a median velocity of 0.01 ± 0.41 µm/s (± IQR, **Fig. 2d** lower panel). The pattern of motility observed with DDB-KIF16B-vesicles (fast unidirectional runs in either direction as well as occasional pauses and directional reversals) closely resembles the behavior observed with native cargoes *in vivo* ^6,7,9,32,33^. Therefore, we conclude that we successfully reconstituted vesicle motility which mimics the transport of intracellular cargoes by multiple opposite-polarity motors.

### Opposing motors do not affect the velocity of the driving motor

Directional reversals of DDB-KIF16B-vesicles are an outcome of the simultaneous presence and activity of DDB and KIF16B on individual vesicles. However, unidirectional runs in either direction can be a result of either: (i) opposite-polarity motors not being present on the vesicles at the same time (i.e. KIF16B is not present during minus-end runs and DDB is not present during plus-end runs) or (ii) opposite-polarity motors being present at the same time but not hindering the driving motors. To distinguish between these two scenarios, we performed a dual-color motility assay where Atto647N-labelled vesicles were incubated with unlabeled DDB and eGFP-labelled KIF16B. We observed fluorescent signals from KIF16B-eGFP on all vesicles moving towards the plus-end. At the same time, we also observed the KIF16-eGFP signal on 76.2% of the vesicles (48 of 63) moving towards the minus-end (**Extended Data Fig. 3a**, blue arrowheads) and on all vesicles exhibiting reversals (**Extended Data Fig. 3a**, magenta arrowheads). This indicates that, for the most part, opposite-polarity motors are simultaneously present on a moving vesicle.

We tested if the transport direction of vesicles could be biased by simply tuning the relative abundance of DDB and KIF16B. To automate the determination of transport direction of vesicles, we developed a segmentation algorithm that parsed the position-time tracks of vesicles into phases of negative runs, positive runs and pauses. (**Fig. 3a**). Tracks were first parsed into runs and pauses by a mean-square displacement method ^34^ followed by determining the slope of each run (Materials and Methods). Runs correspond to phases of directed transport while pauses correspond to either phases of low velocity with no significant net transport (termed stationary pauses; **Extended Data Fig. 3b** upper panel) or diffusive phases characterized by rapid displacements in either direction with no significant net transport (termed diffusive pauses; **Extended Data Fig. 3b** lower panel). Based only on the composition of the runs, individual tracks were classified as minus tracks (composed of only negative runs), plus tracks (composed of only positive runs) and reversal tracks (composed of at least one negative and one positive run). We reconstituted vesicle transport after incubation at a constant DDB concentration (38 nM) with varying KIF16B concentrations (10 – 75 nM) and determined the proportion of minus, plus and reversal tracks (**Fig 3b**). The bulk motor concentration was used as a proxy for the number of vesicle-bound motors. DDB-vesicles (38 nM DDB) and KIF16B-vesicles (25 nM KIF16B) were used as controls. The frequency of minus tracks reduced while that of plus tracks increased with increasing KIF16B concentration. The inversion in the frequency of plus and minus tracks between 10 nM and 75 nM KIF16B shows that the direction of unidirectional motility is sensitive to this concentration range. The frequency of reversal tracks peaked at 25 nM KIF16B (19.4%). The fraction of negative and positive runs (normalized by the total distance travelled) also showed a similar trend: negative runs decreased while positive runs increased with increasing KIF16B concentration (**Extended Data Fig. 3c**). We conclude that vesicles exhibiting directional reversals can be biased to move unidirectionally by increasing the number of motors moving in that particular direction.

**Figure 3.**
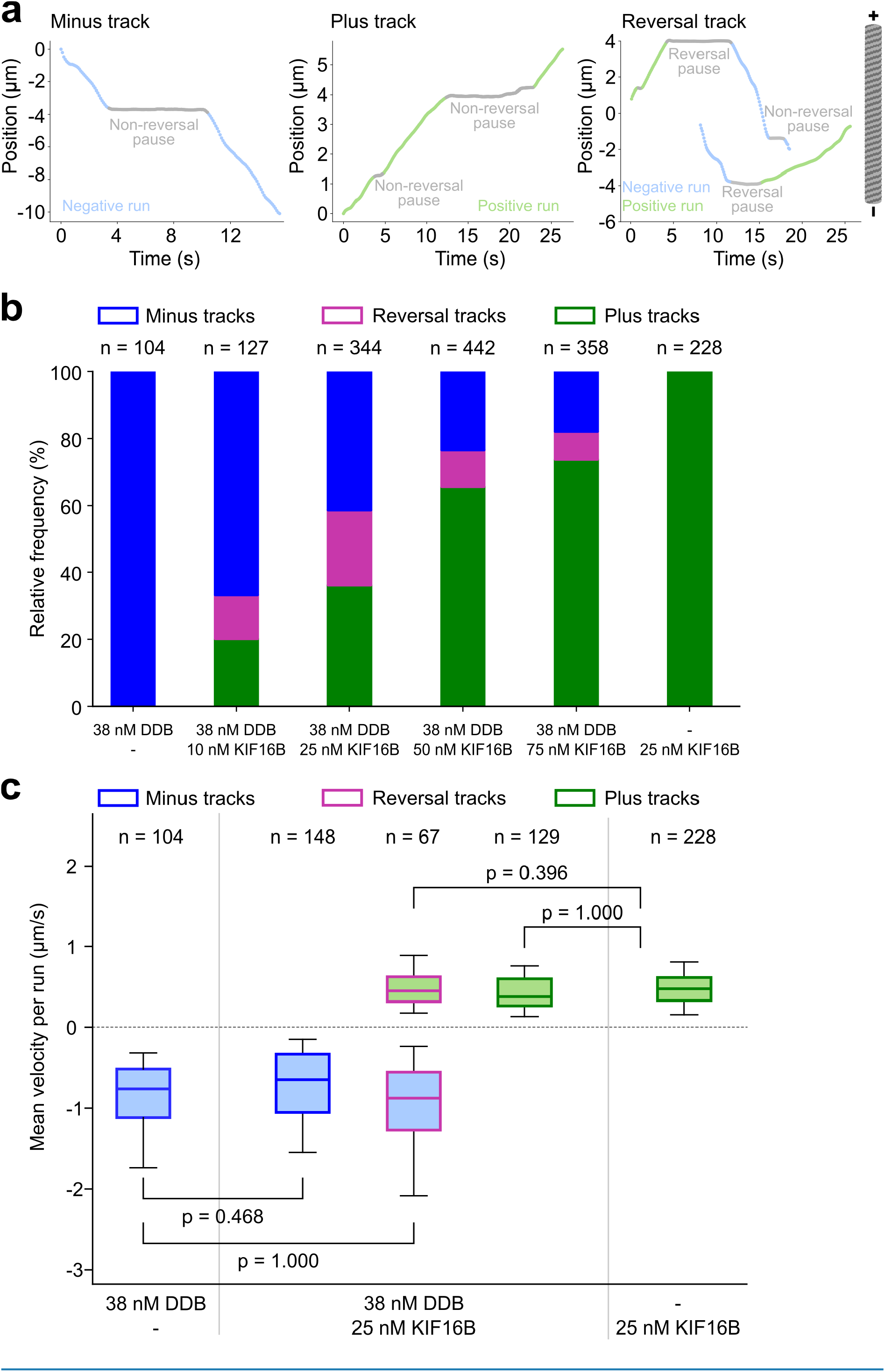
Opposing motors do not affect the velocity of the driving motors. **a)** Position-time tracks of vesicles that were segmented into runs and pauses using a segmentation algorithm based on the mean-square displacement method (see Material and Methods) (Gal et al., 2013). Tracks were classified as minus tracks when they consisted of only negative runs, plus tracks when they consisted of only positive runs and reversal tracks when they consisted of at least one positive and one negative run. Pauses in between the runs were classified as non-reversal pauses when the run direction remained the same and reversal pauses when the run direction reversed after the pause. **b)** Proportion of minus (blue), reversal (magenta) and plus (green) tracks obtained from DDB-KIF16B-vesicles incubated with 38 nM DDB and various concentrations of KIF16B (10, 25, 50, 75 nM). DDB-vesicles and KIF16B-vesicles were used as controls. The data are pooled from two independent experiments. n represents the total number of tracked vesicles for a given condition (data pooled from two independent experiments). **c)** Box plots of mean velocities of negative runs (blue filled boxes) and positive runs (green filled boxes) from minus (blue outline), plus (green outline) and reversal tracks (magenta outline) obtained from DDB-KIF16B-vesicles incubated with 38 nM DDB and 25 nM KIF16B. Negative velocities from minus tracks of DDB-vesicles (38 nM DDB) and positive velocities from plus tracks of KIF16B-vesicles (25 nM KIF16B) are shown as controls. The boxes span 25-75 percentile values, the whiskers span 5-95 percentile values, and the line within the box marks the median. n represents the number of tracked vesicles. Data from two independent experiments performed on two different days were used for the analysis. p-values were computed by comparing the respective samples using an independent t-test with Bonferroni correction.

Although the presence of opposing motors was not sufficient to induce reversals in the transport of all vesicles, it might still alter the characteristics of the runs. Microtubule binding and force generation by opposing motors might cause a reduction in the transport velocity of the driving motors. This kind of interference has been observed previously for artificial assemblies of dynein and kinesin ^19,21,22^ and for microtubules gliding on a lawn of surface-bound dynein and kinesin ^23^. We asked if a similar reduction in velocity also occurs during phases of unidirectional runs for vesicles undergoing reversals. We addressed these possibilities by investigating (i) whether vesicles in our dual-motor vesicle assays were slower than vesicles in our single-motor vesicle assays and (ii) whether the reversing vesicles in our dual-motor vesicle assays were slower than vesicles exhibiting unidirectional motion only.

Towards this end, we focused on vesicles incubated with 38 nM DDB and 25 nM KIF16B as we observed the highest proportion of reversal tracks under this condition. When comparing the negative velocities (**Fig. 3c**, blue filled boxes; **Extended Data Table 2** and **3**) of DDB-vesicles and dual-motor vesicles (from minus tracks and reversal tracks, blue and magenta outlines), we did not observe a considerable reduction in the velocity of the negative runs in the presence of KIF16B. Likewise, positive velocities (**Fig. 3c**, green filled boxes; **Extended Data Table 2** and **3**) of KIF16B-vesicles and dual-motor vesicles (from plus tracks and reversal tracks, green and magenta outlines) were not significantly different. Further supporting these findings, the velocity histogram of dual-motor vesicles, especially the negative and positive tails, resembled a combination of the velocity histograms of DDB-vesicles and KIF16B-vesicles (**Extended Data Fig. 3d)**. Similar results were observed for vesicles incubated with 38 nM DDB and 10, 50 and 75 nM KIF16B (**Extended Data Fig. 3e, Extended Data Table 2** and **3**). Therefore, we conclude that DDB and KIF16B, as opposing motors, do not functionally interfere with the activity of the driving motors during unidirectional runs.

### DDB and KIF16B engage in a tug-of-war during vesicle pausing

While we did not observe a slowdown of vesicles in the dual-motor vesicle assay during unidirectional runs, pausing became the dominant feature in the motility of DDB-KIF16B-vesicles (see major peak around zero in the histogram of instantaneous velocities in **Fig. 2d**). These pauses might indicate periods where DDB and KIF16B simultaneously generate force against each other, i.e., engage in a tug-of-war with no net movement. However, we also observed pauses in single-motor vesicle assays (DDB-vesicles and KIF16B-vesicles), suggesting that not all pauses were a result of a tug-of-war between DDB and KIF16B. Such pauses could arise from a fraction of motors on the vesicles which are inactive (i.e. they interact with the microtubule in a stationary manner) or diffusive (as observed in **Fig. 1b**). Inactive motors comprise motors which are inactive by themselves, pause intermittently between processive runs or are prevented from stepping by lattice defects on the microtubule. Therefore, we investigated if the pauses of dual-motor vesicles were different from the pauses of single-motor vesicles. Focusing again on vesicles incubated with 38 nM DDB and 25 nM KIF16B, we measured the spatial frequency of pauses (number of pauses per micrometer of distance travelled) by calculating the mean of the ratio of the number of pauses to the distance travelled by individual vesicles. We weighted this ratio by the fraction of distance travelled by all vesicles of a given type (i.e. from either minus, plus or reversal tracks; Materials and Methods) as there were many short tracks which did not pause or reached the microtubule ends. Among the dual-motor vesicles, vesicles exhibiting directional reversals had a 2-fold higher spatial pause frequency (0.43 ± 0.04 µm^-1^, weighted mean ± weighted standard error of mean) than unidirectional vesicles (0.20 ± 0.02 µm^-1^, **Fig 4a**). On the other hand, the spatial pause frequency of single-motor vesicles was similar to that of dual-motor vesicles moving unidirectionally only. Given that most unidirectional vesicles in the dual-motor vesicle assay also contain the opposing motor, we conclude that the opposing motor activity was higher on vesicles exhibiting directional reversals as compared to unidirectional vesicles.

**Figure 4.**
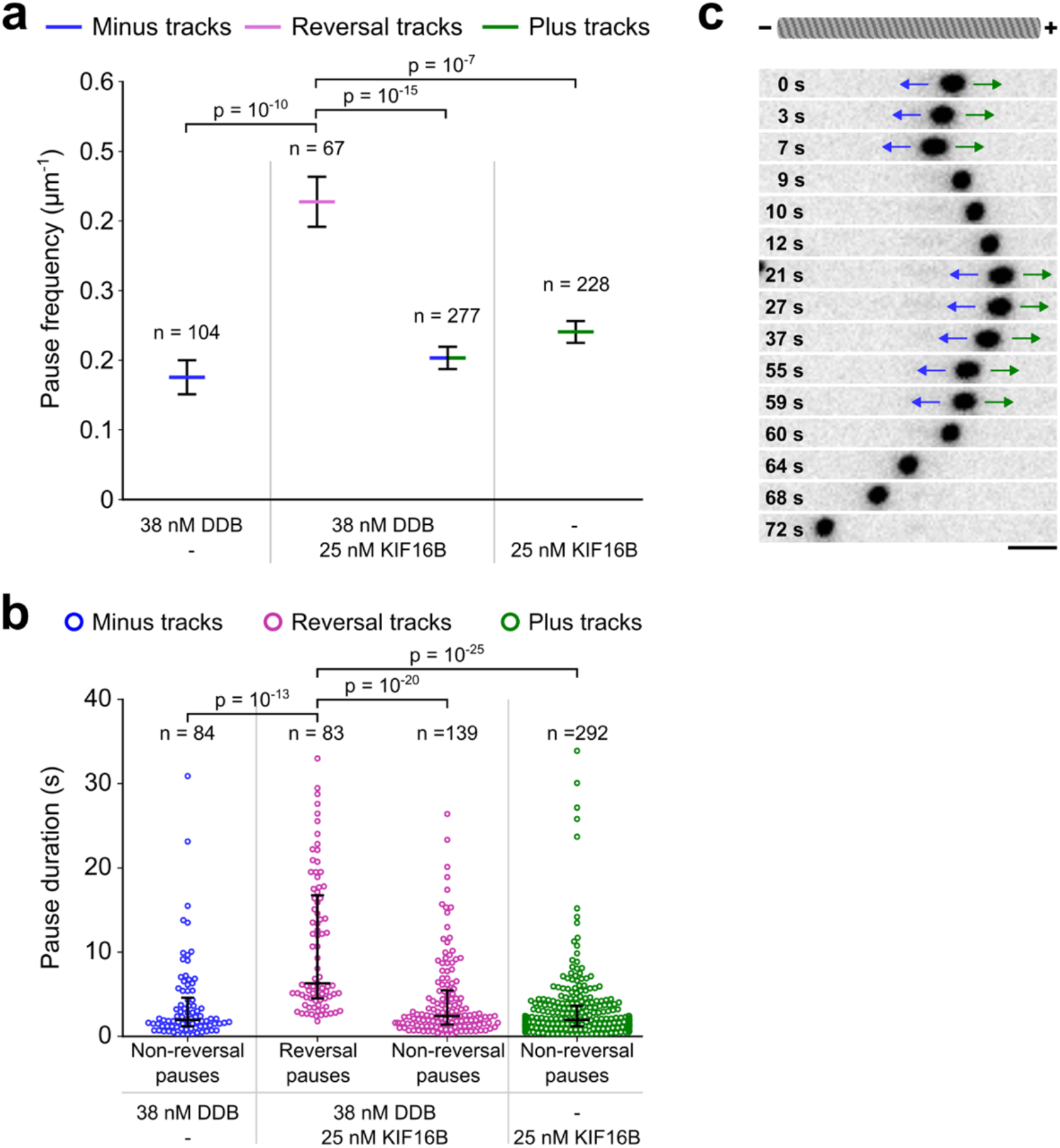
DDB and KIF16B engage in a tug-of-war during vesicle pausing. **a)** Bar plots of spatial pause frequency defined as the number of pauses divided by the distance travelled per track and weighted by the proportion of total distance travelled by vesicles of a given type (minus, plus or reversal tracks) in single- and dual-motor vesicle assays. Spatial pause frequencies of minus and plus tracks from dual-motor vesicles were combined. The height of the bars mark the weighted means and the error bars represent the weighted standard error of the mean (see Materials and Methods). n represents the number of tracks analyzed in each condition. p-values were obtained from two-sample Kolmogorov-Smirnov tests with Bonferroni correction. **b)** Beeswarm plots of pause durations of minus tracks from DDB-vesicles, plus tracks from KIF16B-vesicles and reversal tracks from DDB-KIF16B-vesicles. The central lines mark the median durations while the whiskers span the 25-75 percentile values. p-values were computed by pairwise Mann-Whitney U tests with Bonferroni correction. n represents the number of pauses for a given condition. **c)** Timelapse images of a large vesicle undergoing an elongation along the long axis of a microtubule (not shown). Vesicle elongation coincides with paused states (blue and green arrows at 0-7 s and 21-59 s, where the velocity is slower than 0.2 µm/s) while the vesicle becomes spherical again during fast runs (where the velocity is faster than 1.0 µm/s, at 9-12 s with movement to the plus end and at 60-72 s with movement to the minus end). Scale bar: 2 µm.

Pauses in reversal tracks of dual-motor vesicles can be classified as “reversal pauses” when the transport direction of the vesicle changed after the pause or as “non-reversal pauses” when the run continued in the same direction as before the pause. To investigate if these two types of pauses were characteristically different, we compared their durations with each other and also with pauses of single-motor vesicles. We found that the duration of reversal pauses (6.3 ± 12.3 s, median ± IQR) was significantly higher than the duration of (i) non-reversal pauses (2.4 ± 4.1 s) in the reversal tracks of our dual-motor vesicle assays, (ii) pauses of DDB-vesicles (2.0 ± 3.4 s) and (iii) pauses of KIF16B-vesicles (2.0 ± 2.4 s, **Fig. 4b**). We interpret the longer reversal pause durations to be an outcome of a tug-of-war between DDB and KIF16B, in line with previous reports^19–22^. Non-reversal pauses in the reversal tracks of our dual-motor vesicle assays are a combination of (i) pauses due to a tug-of-war between opposite-polarity motors and (ii) pauses due to the inactivity or intermittent pausing of few motors of the same kind, as these pauses are also observed for single-motor vesicles. However, it is not possible to distinguish between these two possibilities just based on the pause durations. Occasionally, however, we observed elongations of larger vesicles during pausing (**Fig. 4c; Supplementary video 6**). The vesicles elongated towards both directions indicating a tug-of-war, i.e. the simultaneous and independent activity of the associated DDB and KIF16B motors. Moreover, the elongations often occurred during the pauses before the vesicle reversed its direction and disappeared during unidirectional runs. Similar elongations resulting from a tug-of-war between dynein and kinesin have been reported *in vivo* for endosomes of *D. discoideum*^9^.

### Low numbers of attached motors are critical to observe reversals

While previous experimental attempts to reconstitute bidirectional transport *in vitro* resulted in stationary cargoes due to a constant tug-of-war between dynein and kinesin, various mathematical models have been able to recapitulate unidirectional runs and reversals using experimentally determined parameters^9,35,36^. To identify parameters that are key to obtaining reversals and to obtain insights into the dynamics that cause and eventually resolve a tug-of-war, we developed a stochastic stepping model (**Fig. 5a**, Materials and Methods). Briefly, DDB and KIF16B are modelled as Hookean springs on a spherical cargo. As observed in the experiments, the motors are a mixture of active, inactive and diffusive motors. Our cargo is divided into a small attachment area and a large reservoir. Motors present within the attachment area can attach to the microtubule and can generate forces. Upon detachment from the microtubule, motors exchange for new motors (either active, inactive or diffusive; with an individual force-free stepping rate) of the same type (DDB or KIF16B) from the reservoir (see **Extended Data Tables 4 and 5** for the parameters used). This way, we account for the diffusion of motors within the vesicle membrane. The model with either DDB or KIF16B alone is able to recapitulate the experimentally observed instantaneous velocity histograms of single-motor vesicles (**Extended Data Figs. 4a and 4b**).

**Figure 5.**
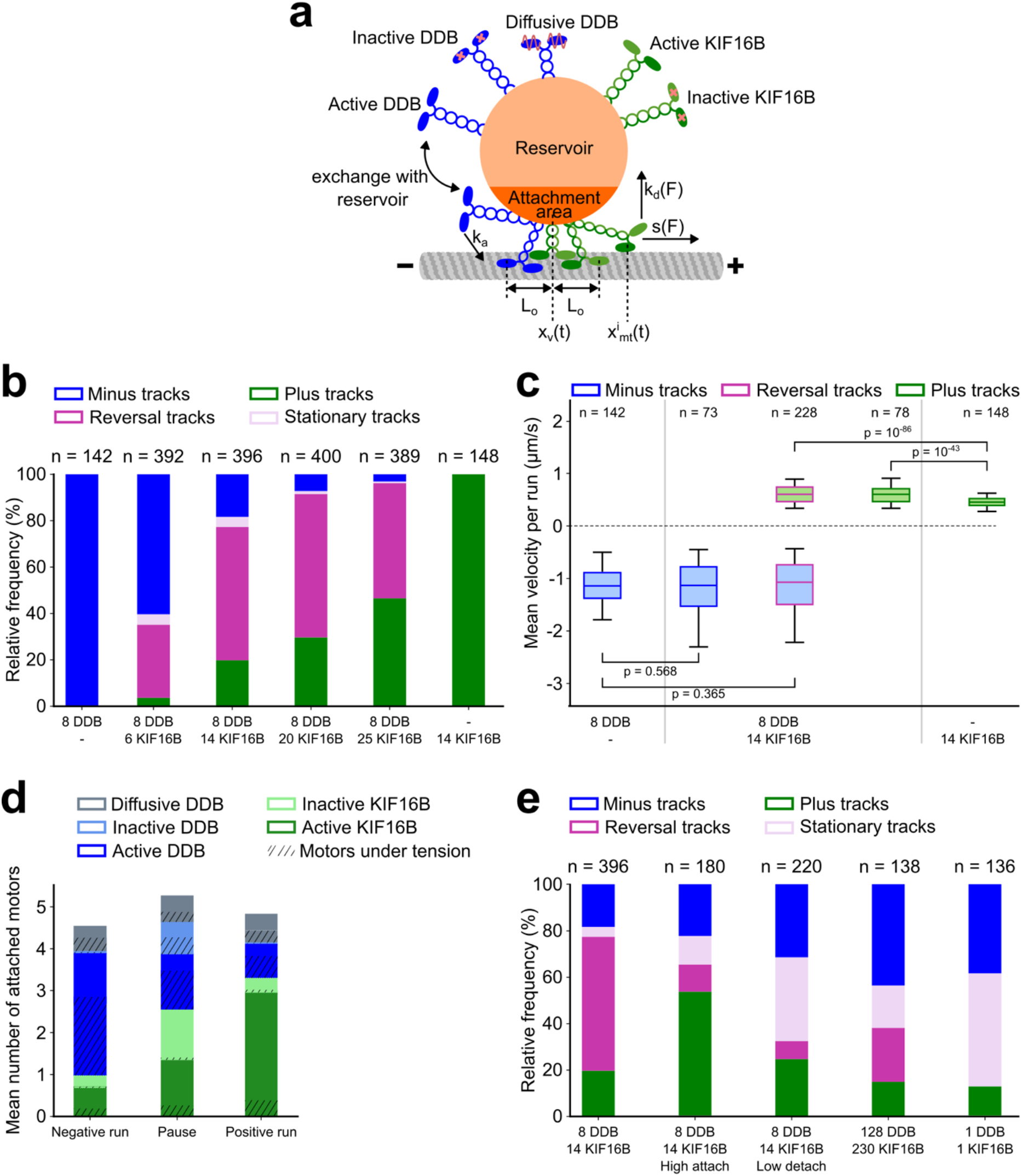
Low numbers of attached motors are critical to observe reversals. **a)** Schematic of the model. Motors bound to the cargo are composed of active KIF16B (green), inactive KIF16B (green with red cross), active DDB (blue), inactive DDB (blue with red cross) and diffusive DDB (blue with red wave). The cargo is divided into an attachment area and a reservoir. Motors in the attachment area can attach the microtubule with a constant attachment rate k_a_. Active DDB and active KIF16B motors, which are attached to the microtubule, step with force dependent stepping rates s(F) to the microtubule minus- and plus-ends, respectively. Attached diffusive DDB motors diffuse in the harmonic potential of their motor springs. All attached motors detach with force-dependent detachment rates k_d_(F). Motors which are stretched outside L_0_ are under tension and exert a force on the cargo (|F| > 0). **b)** Proportions of minus (blue), plus (green), reversal (magenta), and stationary (lilac) tracks obtained from simulations of cargoes with 8 DDB motors and varying numbers of KIF16B motors (6, 14, 20, 25) in the attachment area. Simulations with either 8 DDB (extreme left) or 14 KIF16B motors (extreme right) are shown as controls. n represents the total number of simulated vesicles for a given condition. **c)** Box plots of mean velocities of negative runs (blue filled boxes) and positive runs (green filled boxes) from minus (blue outline), plus (green outline) and reversal tracks (magenta outline) obtained from simulations of cargoes with 8 DDB and 14 KIF16B motors in the attachment area. Negative velocities from minus tracks of DDB-vesicles (8 DDB motors) and positive velocities from plus tracks of KIF16B-vesicles (14 KIF16B motors) are shown as controls. The boxes span 25-75 percentile values, the whiskers span 5-95 percentile values, and the line within the box marks the median. n represents the total number of simulated vesicles for a given condition. p-values were computed from pairwise Mann-Whitney U test (for negative velocities) and pairwise Welch’s test (for positive velocities) with Bonferroni correction to account for multiple comparisons. **d)** Stacked bar plots of mean attached numbers of motors during negative runs (left), pauses (middle) and positive runs (right) from simulations of cargoes with 8 DDB and 14 KIF16B motors in the attachment area. Active KIF16B motors are marked dark green, inactive KIF16B in light green, active DDB in dark blue, inactive DDB in light blue and diffusive DDB in gray. Shaded areas show motors which are under tension. Data from n = 396 simulated cargoes was used to construct this plot. **e)** Proportion of minus (blue), plus (green), reversal (magenta), and stationary (lilac) tracks obtained from simulations of cargoes with (i) 8 DDB and 14 KIF16B motors in the attachment area, (ii) 8 DDB and 14 KIF16B motors with 32-fold higher attachment rates each, (iii) 8 DDB and 14 KIF16B motors with 20-fold lower detachment rates each, (iv) 128 DDB motors and 230 KIF16B motors, and (v) one active DDB motor competing against one active KIF16B motor. n represents the total number of simulated cargoes for a given condition.

We simulated tracks of cargoes with a constant mean number of DDB motors (N_D_ = 8) and varying mean numbers of KIF16B motors (N_K_ = 6, 14, 20, 25) in the attachment area (Materials and Methods). We then segmented these tracks into runs and pauses as described before (Materials and Methods, **Extended Data Fig. 5a**). Similar to our experimental results from dual-motor vesicles, we obtain minus, plus and reversal tracks. The proportion of plus tracks again increases upon increasing the number of KIF16B motors (**Fig. 5b**, compare to **Fig. 3b**). We also identify a small number of stationary tracks where the cargo only paused. Velocities of unidirectional runs are not significantly affected by the presence of opposing motors (**Fig. 5c, Extended Data Fig. 5b, Extended Data Tables 6** and **7**). Moreover, DDB and KIF16B engages in tugs-of-war recapitulating the experimentally observed prolonged durations of reversal pauses compared to non-reversal pauses (**Extended Data Fig. 5c, Extended Data Table 8**). Removing the inactive motors yields similar results (**Extended Data Figs. 6a** and **6b, Extended Data Table 9**) except for the pauses becoming shorter and more diffusive (**Extended Data Fig. 6c)**.

Importantly, our numerical simulations allow us to shed light on the configurations of the motors interacting with the microtubule during the runs and pauses (**Fig. 5d**). During the pauses we observe a high number of attached inactive motors and comparable numbers of attached active KIF16B and active DDB motors. In contrast, unidirectional (positive and negative) runs are characterized by high numbers of active driving motors along with low numbers of inactive and opposing motors. Moreover, we observe that during the runs and pauses only a small fraction of the active attached KIF16B motors are under tension, with the rest forming a pool of untensioned motors. In contrast, a large fraction of the active attached DDB motors are tensioned (**Fig. 5d**). Force balance, e.g. during pausing, is thus maintained by two distinctive mechanisms: Tensioned KIF16B motors exhibit a high force-dependent detachment rate, because only few motors share the load. However, after detachment, untensioned KIF16B motors are readily available to get under tension and take over the load. Tensioned DDB motors, on the other hand, exhibit a low force-dependent detachment rate, because many motors share the load. Together with their lower force-free detachment rate (compared to KIF16B), they thus resist the load for significantly extended time periods.

Utilizing our numerical simulations, we were interested in identifying the conditions that lead to directional reversals. In particular, it is known that the attachment/detachment kinetics and number of motors can be critical to cargo transport, especially during mechanical competition between opposite-polarity motors^37,38^. For example, individual motors rigidly bound to cargo may rapidly reattach (i.e. significantly faster than out of solution) after detachment due to their immediate and retained proximity to the microtubule. However, in case of vesicular cargo, motor diffusion in the membrane is expected to prevent such rapid reattachment. We, therefore, tested if increasing the attachment rate of the motors would reduce the likelihood of directional reversals. We simulated cargo transport with 8 DDB and 14 KIF16B motors with increased attachment rates (32-fold higher for both DDB and KIF16B). A large fraction of tracks then are stationary (**Fig. 5d**). Moreover, the number of attached motors is increased (**Extended Data Figs. 7a**) and the velocities of unidirectional runs in the presence of opposing motors are reduced (**Extended Data Figs. 7b, Extended Data Tables 10** and **11**). We can recapitulate the same effect by either reducing the detachment rate of the motors (20-fold lower for both DDB and KIF16B) or by simply increasing the number of motors bound to the cargo (128 DDB and 230 KIF16B motors) (**Fig. 5e**). When modeling only one KIF16B motor competing against one DDB motor, we observe exclusively stationary or unidirectional tracks, which last only up to a few seconds before detachment of the motor assembly.

## Discussion

Intracellular cargoes undergo directional reversals but transport in either direction is characterized by fast runs^32,33^. Up to now, only non-vesicular assemblies of dynein and kinesin (either as individual motors or as ensembles) have been reconstituted and neither fast unidirectional runs nor directional reversals could be recapitulated in those experiments. The studied assemblies rather exhibited slow motility or highly stable tugs-of-war, characterized by long stationary phases^19–23^. This suggested that dynein and kinesin cannot be active simultaneously on intracellular cargoes^20,36,39^. Simulations that modelled a potential tug-of-war between dynein and kinesin as a stochastic process with unequal load sharing did not fit the experimental data of bidirectionally moving lipid droplets in *Drosophila* embryo^36^. Consequently, dynein and kinesin activities on intracellular cargoes were proposed to be reversibly coordinated by external regulators^10,32,40–43^. These regulators were hypothesized to prevent tugs-of-war by reciprocally activating dynein and kinesin, thereby preventing cargoes from remaining stationary. However, in our *in vitro* experiments with purified DDB motor complexes and KIF16B motors attached to synthetic vesicles we observed transport with similar features as vesicles moving *in vivo*. In particular, our assay recapitulated phases of unidirectional runs and intermittent pauses as well as directional reversals. Velocities of unidirectional runs, both towards the minus- or the plus-end, were not significantly affected by the presence of the opposing motors. Vesicles that reversed their transport direction paused for longer durations than vesicles which paused during unidirectional runs. During pausing, a number of vesicles were observed to elongate along the long axis of the microtubule before reversing their directions. Taken together, our results suggest that the vesicle-bound opposite-polarity motors do not slow down transport during runs, but engage in a tug-of-war during the pauses where stochastic fluctuations in the number of engaged motors can lead to directional reversals without the necessity of external regulators (**Fig. 6**).

**Figure 6.**
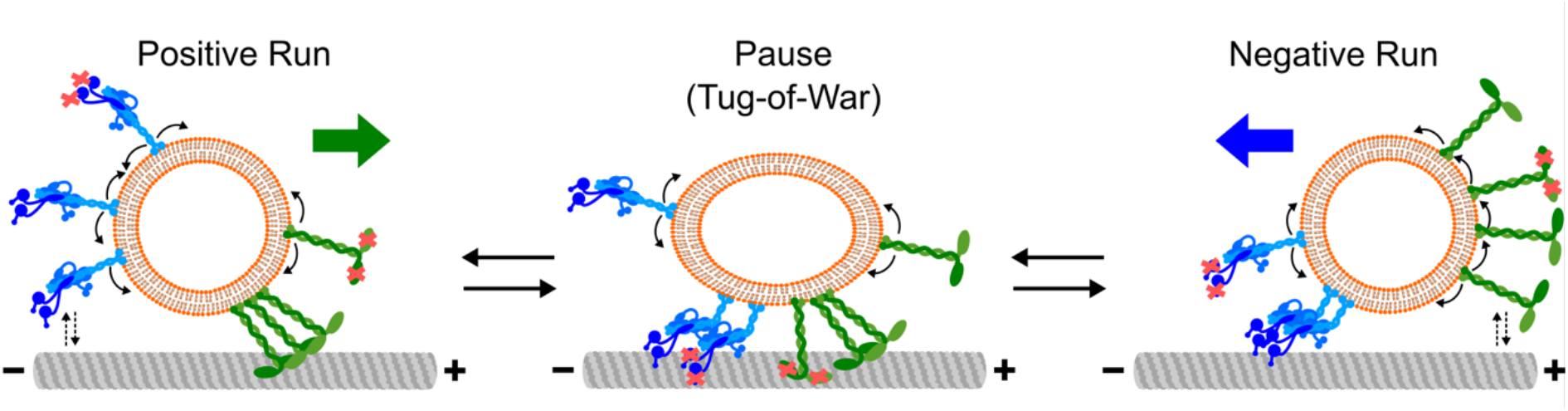
Model for directional reversals of vesicles driven by opposite-polarity motors. Active and inactive (with red cross) DDB and KIF16B motors can diffuse (curved arrows) within the vesicle membrane. A negative or a positive run is initiated by the attachment of active DDB or KIF16B motors to the microtubule, respectively. Only the driving team of motors is attached to the microtubule during the runs. Opposing motors occasionally attach to the microtubule but high load forces by the driving motors cause them to rapidly detach (dashed arrow pairs). Stochastic motor attachment/detachment events result in fluctuations in the number and type of engaged motors. These events include different permutations of (i) attachment of opposite-polarity motors or (ii) attachment of inactive motors of either type or (iii) detachment of driving motors. Such fluctuations can result in a tug-of-war between the opposite-polarity motors, leading to vesicle pausing (with occasional elongation). Likewise, these fluctuations can cause the vesicle to transition from a pause to a run in either direction. A low number of engaged motors is critical for the fluctuations to cause these transitions and to result in directional reversals.

Our numerical simulations allow us to gain information about the motor configurations during bidirectional transport, which is not obtainable from the experimental data. We find that runs are characterized by 2-3 driving active motors and the occasional attachment of opposing and inactive motors (mean number less than 1, **Fig. 5d**). To transition to a pause (characterized typically by a force balance between about 1 active motor of each kind, stabilized by 1-2 inactive motors of either type), 1-2 driving active motors need to detach and 1 opposing as well as 1-2 inactive motors need to attach. A new run in either direction can then be initiated by the detachment of 1 active motor or the attachment of 1-2 active motors (which pull off the inactive and opposing motors). Such changes in the composition of attached motors occur due to stochastic attachment and/or force-dependent detachment events. Thus, in particular at low numbers of attached motors, single-motor detachment and attachment events can already change the state of motility and lead to directional reversals.

When increasing the number of attached motors in our simulations, directional reversals are suppressed. Explaining this observation, we find that transitions between states of motility are more difficult because (i) more motors need to be exchanged and (ii) attachment of additional motors is reduced due to the lack of space on the microtubule. With regard to the latter, we see that even at high numbers of motors bound to the attachment area (e.g.128 DDB together with 230 KIF16B motors) only about 12-14 motors are attached to the microtubule (**Extended Data Figs. 7a**), suggesting that the number of attached motors saturates around this value due to the lack of space on the microtubule. Thus, at high numbers of motors, pauses and runs are stabilized, while directional reversals are suppressed. We note, that high attachment rates or low detachment rates not only increase the number of attached motors, but also favor configurations where almost all available motors are attached to the microtubule. Consequently, only few detached motors are available to change the configuration of the attached motors in terms of directionality and activity. This effect also plays a role at very low numbers of motors, especially for the one-to-one competition between DDB and KIF16B (**Fig. 5e**). Thus, in order to obtain directional reversals, it is required to have (i) a low number of attached motors, such that there is space on the microtubule and that the stochastic attachment and detachment of motors can readily lead to transitions in the state of motility, and (ii) a pool of unattached motors.

In previously performed in vitro assays, the rigidly coupled motors likely exhibited very high attachment rates due to their constant proximity to the microtubule. We hypothesize, that this feature, which also causes a depletion of the pool of unattached motors, has been the reason why no directional reversals have been observed. In contrast, the rapid diffusion of detached motors away from the microtubule on the membrane of vesicular cargo, leads to a reduced (re)attachment rate such that only a small number of motors is attached at any given time. Stochastic motor attachment and/or detachment events can then readily change the state of motility and lead to directional reversals.

A key feature of our assay is the simultaneous but independent activities of the opposite-polarity motors. This was engineered by attaching the DDB and KIF16B to the vesicle with separate binding strategies. However, binding of dynein and kinesin to intracellular cargoes may be interdependent. Recent *in vitro* studies have shown that individual dynein cargo adaptors such as HOOK3^44^ and TRAK2^45^ can scaffold (and activate, in case of TRAK2) dynein and kinesin into a single dynein-kinesin-adaptor complex. Remarkably, unlike previous DNA-based assemblies of single dynein and kinesin molecules^19,21,22^, these complexes exhibited only fast unidirectional transport (either towards the minus- or the plus-end) indicating the exclusive activity of only one type of motor in any given complex. As such, the recruitment of multiple dynein-kinesin-adaptor complexes to a cargo is functionally equivalent to independently attaching dynein and kinesin, as reconstituted in our experiments. The significance of such simultaneous recruitment to, but exclusive activity of dynein and kinesin on, intracellular cargoes remain to be determined.

What determines the transport direction of cargoes? Previous theoretical studies have identified motor properties (stall force, force-dependent detachment rate, etc.) and relative abundance of the motors as determinants of transport direction^9,23,35,36^. Indeed, we observed a strong influence of relative motor concentrations on the transport direction of vesicles (**Fig. 3b**). Estimating the additional influence of external factors on the attachment, detachment and stepping rates of the two motor species will be an intriguing prospect of our combined experimental and theoretical approach in further studies. For example, landing of single molecules of DDB but not KIF1A (a kinesin-3 motor) was shown to be severely affected on MAP9-decorated microtubules^18^. Likewise, tau exerts differential effects on the processivities of dynein and kinesin^1,16^. This was hypothesized to be responsible for the enhanced minus-end motility of phagosomes on tau-decorated microtubules *in vitro*^1^. As we observe that lower active motor numbers are necessary to recapitulate intracellular cargo motility, it will be interesting to investigate the effect of MAPs that limit the number of active motors on cargo transport.

Taken together, incorporation of lipid membranes in *in vitro* motility assays provides an exciting opportunity to study different facets of multi-motor transport, including but not limited to, motor-membrane interactions^24–26,46,47^ and motor-microtubule interactions^48^. In the future, it will be intriguing to extend the presented vesicle assay to systematically analyze the effects of MAPs, tubulin posttranslational-modifications and cargo adaptors on the various features of intracellular cargo transport.

## Acknowledgements

We thank Samara Reck-Peterson (University of California San Diego) for providing the constructs for DIC2-SNAPf and p62-Halo, Marino Zerial (MPI-CBG, Dresden) for providing the constructs for KIF16B, Regis Lemaitre and the Protein Expression, Purification, and Chromatography Facility of MPI-CBG for providing cloning and expression vectors, generating baculovirus and for providing Sf9 and HEK293 cells, Jens Ehrig and the Molecular Imaging and Manipulation Facility of CMCB at Technische Universität Dresden for assistance with microscopy, Corina Bräuer for technical assistance, Veikko Geyer for fruitful discussions, as well as Marino Zerial, Laura Meissner, Stefan Golfier, Jens Ehrig and Aditya Chhatre for comments on the manuscript. We would like to acknowledge funding from the German Research Foundation (SFB1027), the German Federal Ministry of Education and Research (OptiZeD 03Z22E511) and the Boehringer Ingelheim Fonds (PhD stipend to A. I. D’Souza).

## Author contributions

A.I.D., R.G. and S.D. conceptualized and designed the experimental research; G.A.M. and L.S. conceptualized theoretical modeling and simulations; A.I.D. and R.G. generated the protein constructs and performed the experiments; G.A.M. performed the theoretical modeling and simulations, A.I.D., R.G., G.A.M. analyzed the data; all authors discussed the data; A.I.D., R.G., G.A.M. and S.D. wrote the paper.

