## Extended Data and Materials and Methods for "Vesicles driven by dynein and kinesin exhibit directional reversals without external regulators"

### Extended Data Figures

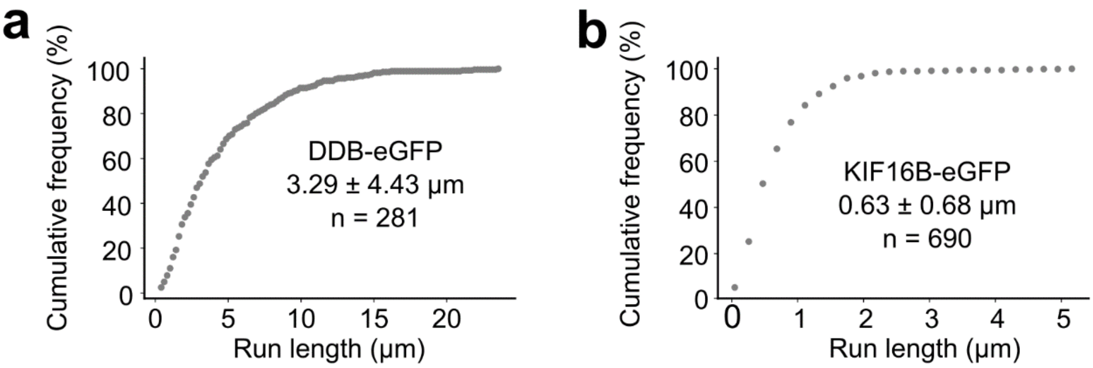


**Figure 1 | Purified DDB-eGFP complexes and KIF16B-eGFP are processive *in vitro*.** Cumulative distribution frequency of run length of **a)** DDB-eGFP and **b)** KIF16B-eGFP. Numerical values are reported as median ± IQR. n represents the number of single-molecules/complexes.

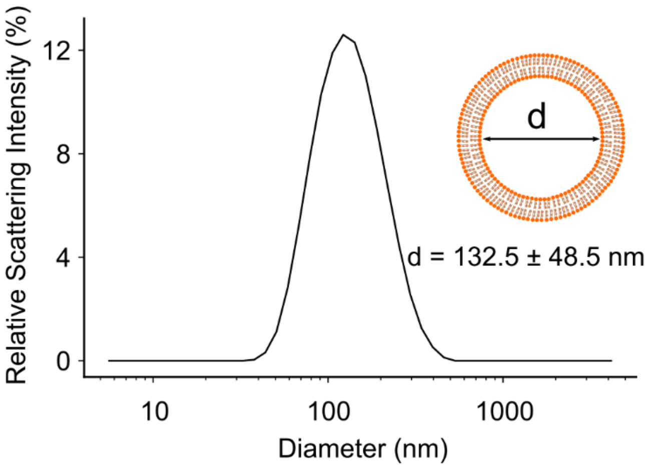


**Figure 2 | Uniformely sized vesicles are obtained by extrusion.** Size distribution of large unilamellar vesicles obtained by dynamic light scattering (Zetasizer, Malvern). The size histogram was obtained from a single DLS run. d is reported as mean ± standard deviation

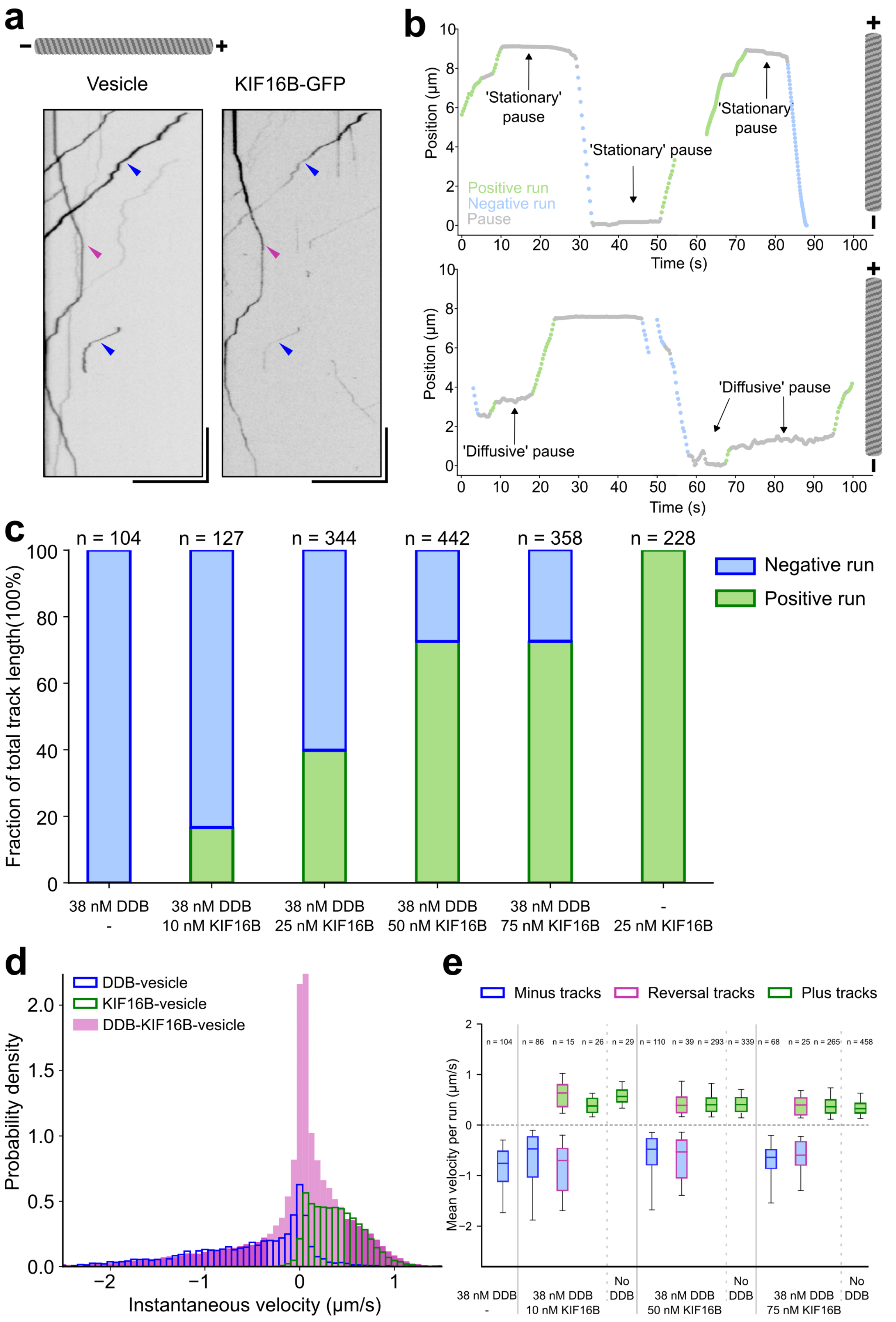


**Figure 3 | Opposing motors do not affect the velocity of the driving motor. a)**

Kymographs of atto647N-labelled vesicles (left panel) incubated with unlabelled DDB and KIF16B-eGFP (right panel) showing colocalizations between KIF16B-eGFP and a reversing vesicle (magenta arrowhead) and a minus-end moving vesicle (blue arrowhead). Scale bars: vertical 20s, horizontal 23 µm. **b)** Position-time tracks of DDB-KIF16B-vesicles with stationary pauses (upper panel) and diffusive pauses (lower panel). **c)** Proportion of negative and positive runs obtained from DDB-KIF16B-vesicles incubated with 38 nM DDB and various concentrations of KIF16B (10, 25, 50, 75 nM). DDB-vesicles and KIF16B-vesicles were used as controls. n represents the total number of tracked vesicles for a given condition (data pooled from two independent experiments). **d)** Probability densities of instantaneous velocity histograms of DDB-vesicles (Fig. 2B) and KIF16B-vesicles (Fig. 2C) were uniformly scaled (factor of 0.355) and overlaid onto the instantaneous velocity histogram of DDB-KIF16B-vesicles. Note that the shape of the high velocity tails (dark magenta) matches between the dual- and single-motor vesicles. **e)** Comparitive mean velocity box plot of negative and positive runs obtained from dual-motor vesicles (38 nM DDB and 10, 50 and 75 nM KIF16B) and single motor vesicles. Numerical values and statistical comparisons are presented In **Extended Data Tables 2** and **3**.

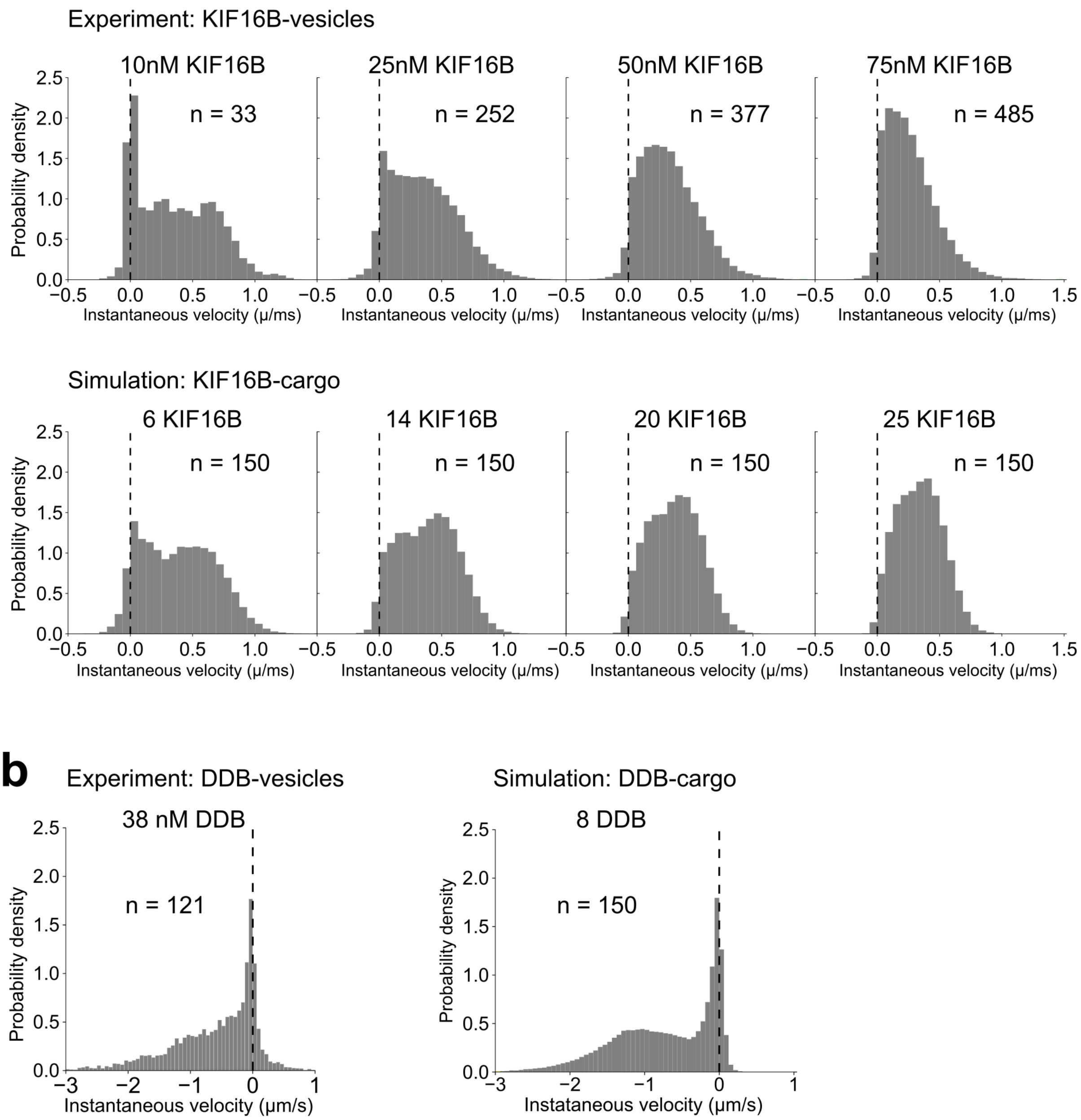


**Figure 4 | Stochastic numerical simulation can recapitulate instantenous velocity profiles of single-motor vesicles. A)** Instantaneous velocity histograms of vesicles incubated with 10, 25, 50 and 75 nM KIF16B (upper panel) and of cargoes simulated with 6, 14, 20 and 25 KIF16B motors in the attachment area (lower panel). **B)** Instantaneous velocity histograms of vesicles incubated with 38 nM DDB (left) and of cargoes simulated with 8 DDB motors in the attachment area (right). n represents the number of motor-bound vesicles/cargoes used to construct the histograms.

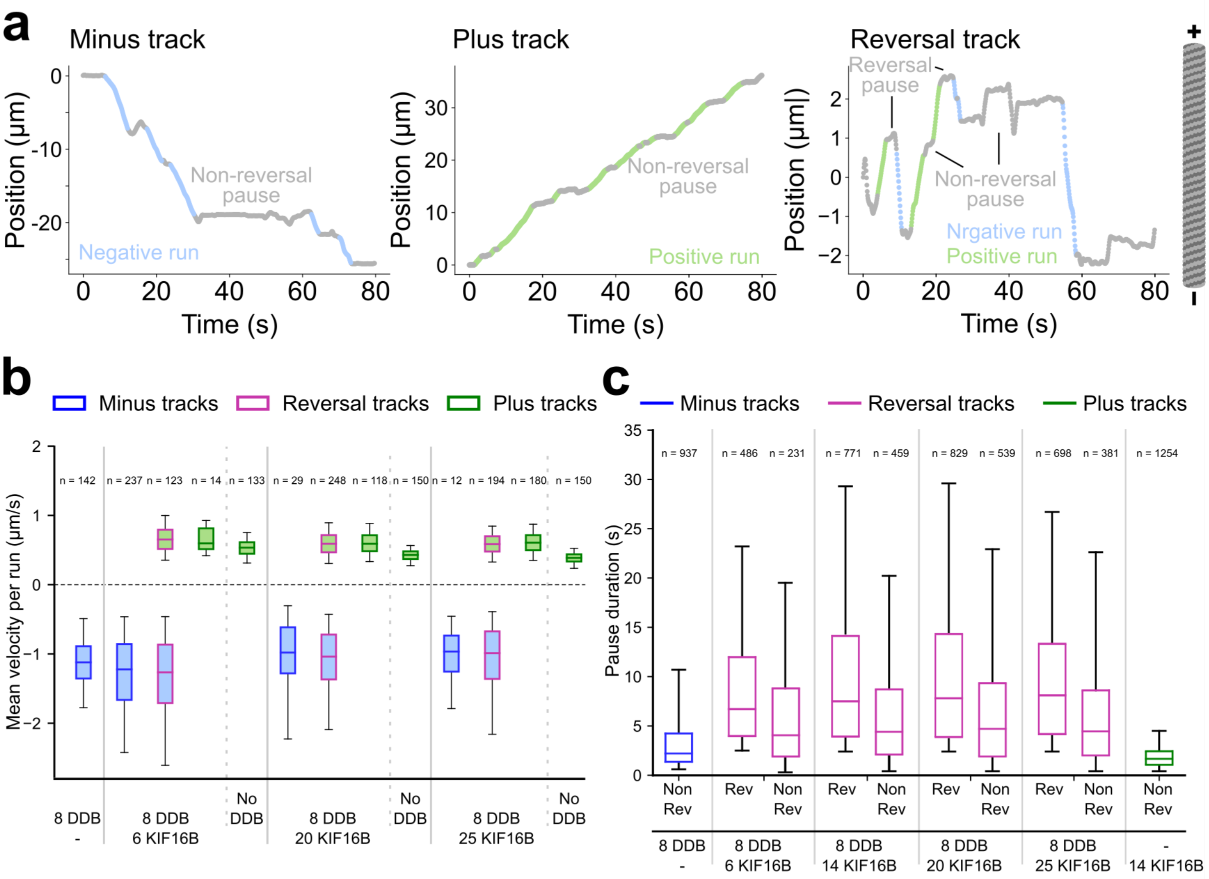


**Figure 5 | Simulated assemblies of dual-motor cargoes recapitulate the characteristics of runs and pauses of dual-motor vesicles. A)** Segmented minus tracks (left), plus tracks (middle), and reversal tracks (right) obtained from simulation of dual-motor cargoes (8 DDB and 14 KIF16B motors) and single-motor cargoes. **B)** Comparitive mean velocity box plot of negative and positive runs obtained from simulation of dual-motor (8 DDB and 6, 20 and 25 KIF16B motors and single motor cargoes. Numerical values and statistical comparison is presented in **Extended Data Tables 6** and **7**. **c)** Comparitive pause duration box plots of reversal and non-reversal pauses obtained from simulation of dual-motor (8 DDB and 6, 14, 20 and 25 KIF16B motors) and single-motor cargoes. Statistical comparisons are presented in **Extended Data Table 8**.

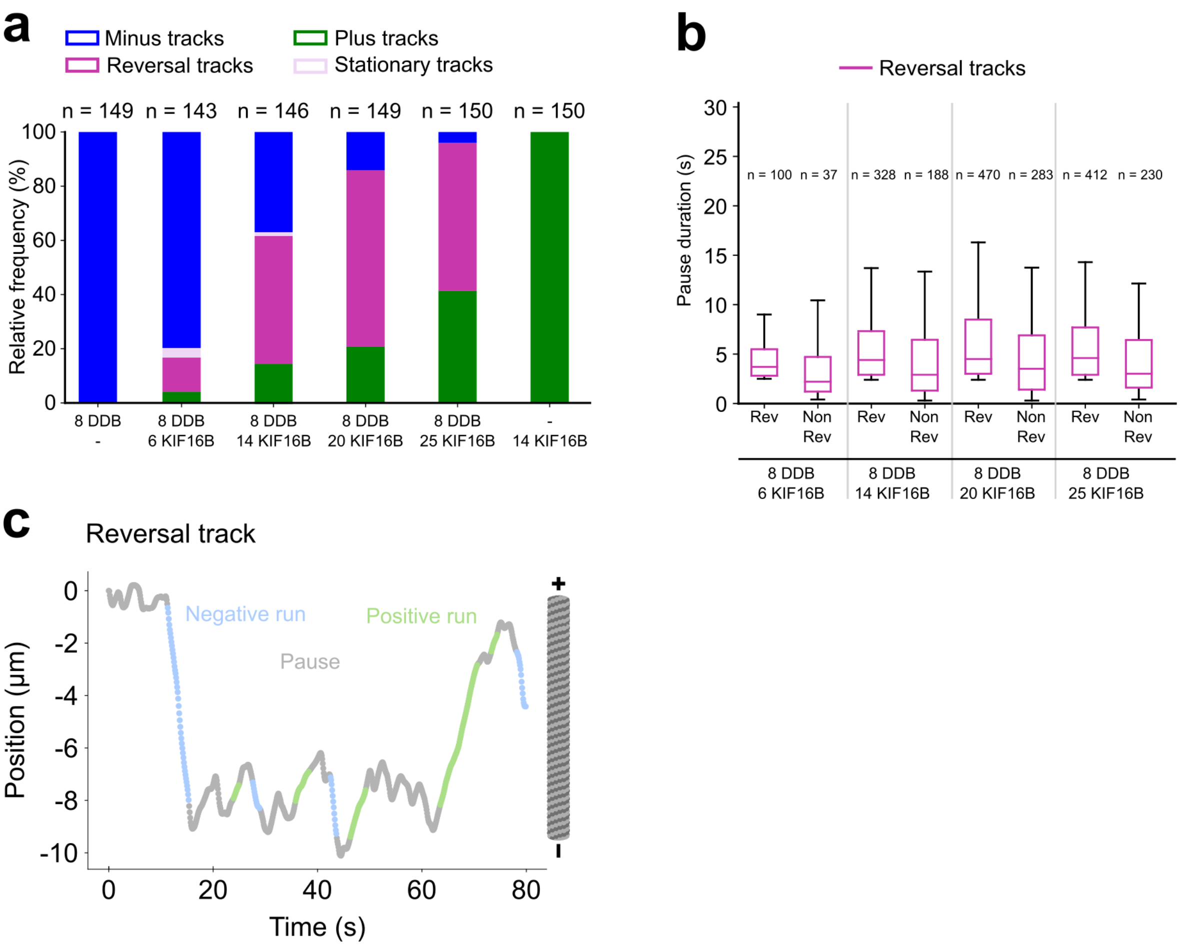


**Figure 6 | Inactive motors affects the dynamics and duration of pauses. a)** Proportions of minus (blue), plus (green), reversal (magenta), and stationary (lilac) tracks obtained from the simulated data. Tracks are obtained from simulations of cargoes with 8 DDB motors and varying numbers of KIF16B motors (6, 14, 20, 25). Simulations with either 8 DDB (extreme left) or 14 KIF16B motors (extreme right) are shown as controls. n represents the total number of simulated cargoes for a given condition. **b)** Comparitive pause duration box plots of reversal (rev) and non-reversal (non-rev) pauses obtained from simulation of dual-motor and single-motor cargoes. Statistical comparisons are presented in **Extended Data Table 9**. **c)** A representative trace of a dual-motor cargo exhibiting directional reversal. The paused phase resemble a diffusive state. All results given in figs. a)-c) are obtained without any inactive motors in the attachment area.

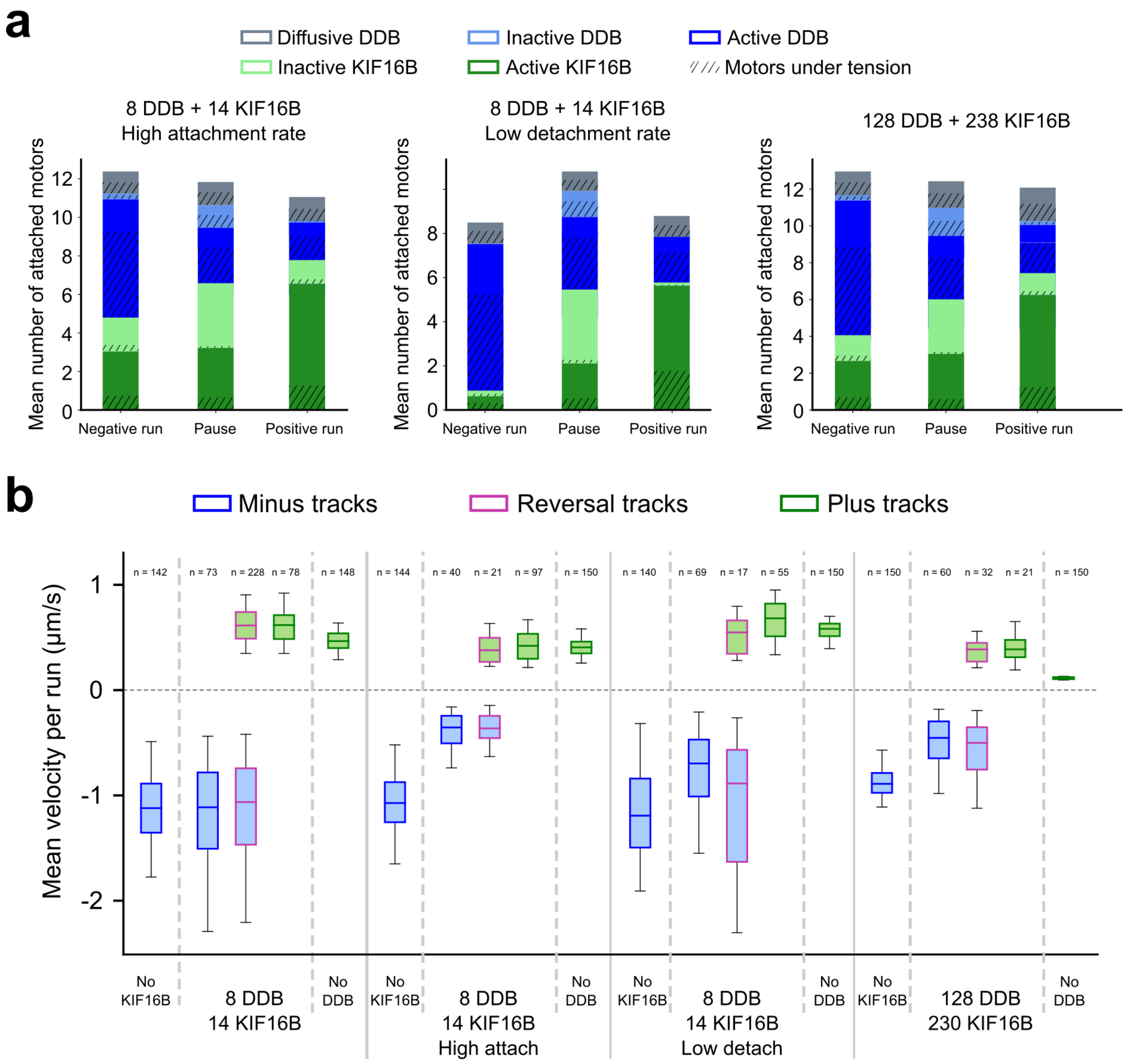


**Figure 7 | Increasing the number of attached motors affects characteristics of dual-motor cargo transport. a)** Stacked bar histograms of mean number of motors attached to the microtubule during negative runs, positive runs and paused phases when dual-motor cargoes were simulated with higher attachment rate motors (left, 20 $s^{-1}$ vs 0.625 $s^{-1}$ for DDB, 40 $s^{-1}$ vs 1.25 $s^{-1}$ for KIF16B), low detachment rate motors (middle, 0.022 $s^{-1}$ vs 0.44 $s^{-1}$ for active DDB, 0.009 $s^{-1}$ vs 0.18 $s^{-1}$ for inactive and diffusive DDB, 0.0635 $s^{-1}$ vs 1.27 $s^{-1}$ for active KIF16B, 0.035 $s^{-1}$ vs 0.7 $s^{-1}$ for inactive KIF16B) and high number of motors (right, 128 DDB and 230 KIF16B). Clear boxes represent untensioned motors while shaded boxes represent motors under tension. **b)** Comparitive mean velocity box plots of negative and positive runs of dual-motor cargoes simulated with altered motor attachment/detachment kinetics and numbers. Numerical values and statistical comparisons are presented in **Extended Data Tables 10** and **11** respectively.

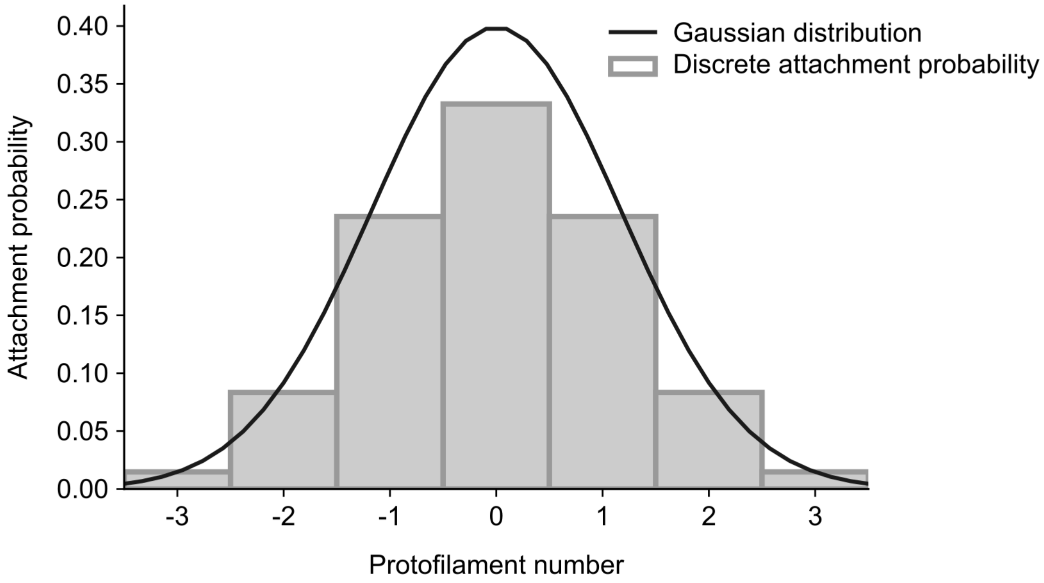


**Figure 8 | Gaussian distributed attachment to protofilaments**

Bar plot shows the attachment probability of either motor type distributed over the microtubule protofilaments. Attachment probability is the highest for the central protofilament (number zero) and reduces in a Gaussian manner for protofilaments away from the central protofilament. The corresponding Gaussian distribution (mean zero and standard deviation one) is shown as solid line. 7 out of 14 protofilaments of a microtubule are accessible to the motors.

### Extended Data Table

| Mol (%) | Short-form | Proper name | Function |
|---|---|---|---|
| 74 | DOPC | 1,2-dioleoyl-sn-glycero-3-phosphocholine | Structural phospholipid |
| 20 | DOPE | 1,2-dioleoyl-sn-glycero-3-phosphoethanolamine | Enhance KIF16B binding |
| 3 | PI(3)P | 1,2-dioleoyl-sn-glycero-3-phospho-(1'-myo-inositol-3'-phosphate) | KIF16B binding |
| 3 | DGS- NTA(Ni) | 1,2-dioleoyl-sn-glycero-3-[(N-(5-amino-1-carboxypentyl)iminodiacetic acid)succinyl] (nickel salt) | DDB binding |
| 0.01 | DOPE-Atto647N | 1,2-dioleoyl-sn-glycero-3-phosphoethanolamine labelled with Atto647N | Fluorescent marker |

**Table1 | Phospholipid composition of vesicles**. Phospholipids were mixed in the indicated molar ratios in chloroform, dried first under a stream of nitrogen gas and then in vacuum for 3-4 hours. Multilamellar vesicles (MLVs) were synthesized by rehydrating the lipid film in dynein buffer B (25 mM HEPES pH 7.4, 50 mM Potassium acetate, 2 mM magnesium acetate) supplemented with 5 % (w/v) sucrose for 30 min at 37 °C (with periodic agitation). An aliquot of MLVs (total lipid mass of 50 μg) was used to synthesize LUVs as outlined in the main text (see also Materials and Methods).

| Motor composition | Direction | Velocity (µm/s) (median ± IQR) | Number of vesicles |
|---|---|---|---|
| 38 nM DDB | Minus | -0.76 ± 0.60 | 104 |
| 38 nM DDB + 10 nM KIF16B | Minus | -0.47 ± 0.79 | 86 |
| | Reversal-Minus | -0.71 ± 0.83 | 15 |
| | Reversal-Plus | 0.63 ± 0.43 | 15 |
| | Plus | 0.38 ± 0.28 | 26 |
| 10 nM KIF16B | Plus | 0.56 ± 0.23 | 29 |
| 38 nM DDB + 25 nM KIF16B | Minus | -0.67 ± 0.73 | 148 |
| | Reversal-Minus | -0.80 ± 0.62 | 67 |
| | Reversal-Plus | 0.45 ± 0.32 | 67 |
| | Plus | 0.38 ± 0.34 | 129 |
| 25 nM KIF16B | Plus | 0.43 ± 0.31 | 228 |
| 38 nM DDB + 50 nM KIF16B | Minus | -0.48 ± 0.52 | 110 |
| | Reversal-Minus | -0.53 ± 0.75 | 39 |
| | Reversal-Plus | 0.39 ± 0.31 | 39 |
| | Plus | 0.40 ± 0.27 | 293 |
| 50 nM KIF16B | Plus | 0.40 ± 0.27 | 339 |
| 38 nM DDB + 75 nM KIF16B | Minus | -0.64 ± 0.36 | 68 |
| | Reversal-Minus | -0.6 ± 0.46 | 25 |
| | Reversal-Plus | 0.39 ± 0.36 | 25 |
| | Plus | 0.36 ± 0.26 | 265 |
| 75 nM KIF16B | Plus | 0.32 ± 0.19 | 458 |

**Table 2 | Median ± IQR of mean instantaneous velocities of experimental vesicles.** Velocities of minus-end directed runs (blue background) and plus-end directed runs (green background) of single- and dual-motor vesicles.

<table>
<tr><th>Motor composition and test</th><th colspan="4">p-values with Bonferroni correction (3 comparisons)<br>- Mean velocities of negative runs-</th></tr>
<tr><td rowspan="4">38 nM DDB +<br>10 nM KIF16B<br>Mann-Whitney U test</td><td></td><td>Only DDB</td><td>Minus</td><td>Reversal-Minus</td></tr>
<tr><td>Only DDB</td><td>1</td><td>0.031</td><td>1</td></tr>
<tr><td>Minus</td><td>0.031</td><td>1</td><td>0.471</td></tr>
<tr><td>Reversal-Minus</td><td>1</td><td>0.471</td><td>1</td></tr>
<tr><td rowspan="4">38 nM DDB +<br>25 nM KIF16B<br>Ind. t-test</td><td></td><td>Only DDB</td><td>Minus</td><td>Reversal-Minus</td></tr>
<tr><td>Only DDB</td><td>1</td><td>0.4681</td><td>1</td></tr>
<tr><td>Minus</td><td>0.4681</td><td>1</td><td>0.1336</td></tr>
<tr><td>Reversal-Minus</td><td>1</td><td>0.1336</td><td>1</td></tr>
<tr><td rowspan="4">38 nM DDB +<br>50 nM KIF16B<br>Mann-Whitney U test</td><td></td><td>Only DDB</td><td>Minus</td><td>Reversal-Minus</td></tr>
<tr><td>Only DDB</td><td>1</td><td>0.00046</td><td>0.0139</td></tr>
<tr><td>Minus</td><td>0.00046</td><td>1</td><td>1</td></tr>
<tr><td>Reversal-Minus</td><td>0.0139</td><td>1</td><td>1</td></tr>
<tr><td rowspan="4">38 nM DDB +<br>75 nM KIF16B<br>Mann-Whitney U test</td><td></td><td>Only DDB</td><td>Minus</td><td>Reversal-Minus</td></tr>
<tr><td>Only DDB</td><td>1</td><td>0.321</td><td>0.046</td></tr>
<tr><td>Minus</td><td>0.321</td><td>1</td><td>0.7738</td></tr>
<tr><td>Reversal-Minus</td><td>0.046</td><td>0.7738</td><td>1</td></tr>
<tr><th>Motor composition and test</th><th colspan="4">p-values with Bonferroni correction (3 comparisons)<br>- Mean velocities of positive runs-</th></tr>
<tr><td rowspan="4">38 nM DDB +<br>10 nM KIF16B<br>Welch's test</td><td></td><td>Reversal-Plus</td><td>Plus</td><td>Only KIF16B</td></tr>
<tr><td>Reversal-Plus</td><td>1</td><td>0.036</td><td>1</td></tr>
<tr><td>Plus</td><td>0.036</td><td>1</td><td>0.01897</td></tr>
<tr><td>Only KIF16B</td><td>1</td><td>0.01897</td><td>1</td></tr>
<tr><td rowspan="4">38 nM DDB +<br>25 nM KIF16B<br>Ind. t-test</td><td></td><td>Reversal-Plus</td><td>Plus</td><td>Only KIF16B</td></tr>
<tr><td>Reversal-Plus</td><td>1</td><td>0.446</td><td>0.396</td></tr>
<tr><td>Plus</td><td>0.446</td><td>1</td><td>1</td></tr>
<tr><td>Only KIF16B</td><td>0.396</td><td>1</td><td>1</td></tr>
<tr><td rowspan="4">38 nM DDB +<br>50 nM KIF16B<br>Mann-Whitney U test</td><td></td><td>Reversal-Plus</td><td>Plus</td><td>Only KIF16B</td></tr>
<tr><td>Reversal-Plus</td><td>1</td><td>1</td><td>1</td></tr>
<tr><td>Plus</td><td>1</td><td>1</td><td>1</td></tr>
<tr><td>Only KIF16B</td><td>1</td><td>1</td><td>1</td></tr>
<tr><td rowspan="4">38 nM DDB +<br>75 nM KIF16B<br>Mann-Whitney U test</td><td></td><td>Reversal-Plus</td><td>Plus</td><td>Only KIF16B</td></tr>
<tr><td>Reversal-Plus</td><td>1</td><td>1</td><td>0.993</td></tr>
<tr><td>Plus</td><td>1</td><td>1</td><td>0.0276</td></tr>
<tr><td>Only KIF16B</td><td>0.993</td><td>0.0276</td><td>1</td></tr>
</table>

**Table 3 | Statistical comparisons of experimental negative and positive velocities of all dual-motor vesicles.** Supplementary to Fig. 3c and Extended Data Fig. 3e

| Parameter | Value | Discussion and references |
| --- | --- | --- |
| Attachment rate | 2.5 $s^{-1}$ (0.625 $s^{-1}$) | Standard (in the presence of KIF16B)[1,2] |
| Force-free detachment rate, active motors | 0.44 $s^{-1}$ | Obtained from experiments (**Fig. 1** and **Extended Data Figs. 1**) |
| Force-free detachment rate, inactive motors | 0.18 $s^{-1}$ | Obtained from comparing simulated and experimental DDB-vesicle instantaneous velocities. |
| Stall force | 4 pN | [3] |
| Detachment force (exponential) | 3 pN | [4] |
| Force-free velocity at 2.5 mM ATP | Instantaneous velocity distribution of single DDB (ignoring positive velocities) | Experiment (**Fig. 1)** |
| Backward velocity | 6 nm/s | Same order of magnitude as [5] |
| Diffusion rate of diffusive DDB motors per 8 nm | 250 $s^{-1}$ | [6] |
| Stiffness | 0.065 pN.$nm^{-1}$ | [7] |
| Untensioned length | 30 nm | Same order of magnitude as [8] |
| Motor radius on the microtubule | 24 nm | Approximated from EM images of [9,10] |

**Table 4 | DDB parameters used in numerical simulations**

| Parameter | Value | Discussion and references |
|---|---|---|
| Attachment rate | 5 $s^{-1}$ (1.25 $s^{-1}$) | Standard (in the presence of DDB)[1,2] |
| Force-free detachment rate | 1.27 $s^{-1}$ | Obtained from experiments (**Fig. 1** and **Extended Data Figs. 1**). |
| Force-free detachment rate for inactive motors | 0.70 $s^{-1}$ | Obtained from comparing simulated and experimental KIF16B-vesicle instantaneous velocities. |
| Detachment force | 2 pN | Same order of magnitude as [11] |
| Stall force | 6 pN | [11] |
| Force-free velocity at 2.5 mM ATP | Instantaneous velocity distribution of single KIF16B motors (ignoring negative velocities) | Experiment (**Fig. 1)** |
| Backward velocity | 6 nm/s | Same order of magnitude as [5] |
| Stiffness | 0.3 pN.$nm^{-1}$ | [7] |
| Untensioned length | 70 nm | Same order of magnitude as [12,13] |
| Motor radius on the microtubule | 4 nm | Same order of magnitude as [14] |

**Table 5 | KIF16B parameters used in numerical simulations**

| Motor composition | Direction | Velocity (µm/s) (median ± IQR) | Number of cargoes |
|---|---|---|---|
| 8 DDB | Minus | -1.12 ± 0.46 | 142 |
| 8 DDB + 6 KIF16B | Minus | -1.22 ± 0.81 | 237 |
| | Reversal-Minus | -1.27 ± 0.84 | 123 |
| | Reversal-Plus | 0.65 ± 0.28 | 123 |
| | Plus | 0.6 ± 0.3 | 14 |
| 6 KIF16B | Plus | 0.53 ± 0.16 | 133 |
| 8 DDB + 14 KIF16B | Minus | -1.11 ± 0.72 | 73 |
| | Reversal-Minus | -1.1 ± 0.73 | 228 |
| | Reversal-Plus | 0.61 ± 0.25 | 228 |
| | Plus | 0.62 ± 0.23 | 78 |
| 14 KIF16B | Plus | 0.47 ± 0.14 | 148 |
| 8 DDB + 20 KIF16B | Minus | -0.98 ± 0.66 | 29 |
| | Reversal-Minus | -1.04 ± 0.65 | 248 |
| | Reversal-Plus | 0.59 ± 0.25 | 248 |
| | Plus | 0.59 ± 0.23 | 118 |
| 20 KIF16B | Plus | 0.43 ± 0.11 | 150 |
| 8 DDB + 25 KIF16B | Minus | -0.97 ± 0.52 | 12 |
| | Reversal-Minus | -0.99 ± 0.69 | 194 |
| | Reversal-Plus | 0.58 ± 0.22 | 194 |
| | Plus | 0.61 ± 0.22 | 180 |
| 25 KIF16B | Plus | 0.39 ± 0.10 | 150 |

**Table 6 | Median ± IQR of means of instantaneous velocities per run of simulated cargoes.** Velocities of minus-end directed runs (blue background) and plus-end directed runs (green background) of simulated single- and dual-motor cargoes. Supplementary to Fig. 5c and Extended Data Fig. 5b

| Motor composition and test | p-values with Bonferroni correction (3 comparisons) - Mean velocities of negative runs- | | | |
|---|---|---|---|---|
| 8 DDB + 6 KIF16B **Mann-Whitney-U test** | | **Only DDB** | **Minus** | **Reversal-Minus** |
| | **Only DDB** | 1 | $10^{-8}$ | $10^{-9}$ |
| | **Minus** | $10^{-8}$ | 1 | 0.33789 |
| | **Reversal-Minus** | $10^{-9}$ | 0.33789 | 1 |
| 8 DDB + 14 KIF16B **Mann-Whitney-U Test** | | **Only DDB** | **Minus** | **Reversal-Minus** |
| | **Only DDB** | 1 | 1 | 0.364515 |
| | **Minus** | 1 | 1 | 0.567571 |
| | **Reversal-Minus** | 0.364515 | 0.567571 | 1 |
| 8 DDB + 20 KIF16B **Mann-Whitney-U Test** | | **Only DDB** | **Minus** | **Reversal-Minus** |
| | **Only DDB** | 1 | 0.00577 | 0.00169 |
| | **Minus** | 0.00577 | 1 | 0.61937 |
| | **Reversal-Minus** | 0.00169 | 0.61937 | 1 |
| 8 DDB + 25 KIF16B **Mann-Whitney-U Test** | | **Only DDB** | **Minus** | **Reversal-Minus** |
| | **Only DDB** | 1 | 0.03017 | $10^{-5}$ |
| | **Minus** | 0.03017 | 1 | 1 |
| | **Reversal-Minus** | $10^{-5}$ | 1 | 1 |
| **Motor composition and test** | **p-values with Bonferroni correction (3 comparisons) - Mean velocities of positive runs-** | | | |
| 8 DDB + 6 KIF16B **Mann-Whitney U test** | | **Reversal-Plus** | **Plus** | **Only KIF16B** |
| | **Reversal-Plus** | 1 | 1 | $10^{-19}$ |
| | **Plus** | 1 | 1 | 0.017 |
| | **Only KIF16B** | $10^{-19}$ | 0.017 | 1 |
| 8 DDB + 14 KIF16B **Welch's test** | | **Reversal-Plus** | **Plus** | **Only KIF16B** |
| | **Reversal-Plus** | 1 | 1 | $10^{-86}$ |
| | **Plus** | 1 | 1 | $10^{-43}$ |
| | **Only KIF16B** | $10^{-86}$ | $10^{-43}$ | 1 |
| 8 DDB + 20 KIF16B **Welch's test** | | **Reversal-Plus** | **Plus** | **Only KIF16B** |
| | **Reversal-Plus** | 1 | 1 | $10^{-136}$ |
| | **Plus** | 1 | 1 | $10^{-114}$ |
| | **Only KIF16B** | $10^{-136}$ | $10^{-114}$ | 1 |
| 8 DDB + 25 KIF16B **Mann-Whitney U test** | | **Reversal-Plus** | **Plus** | **Only KIF16B** |
| | **Reversal-Plus** | 1 | 0.021 | $10^{-183}$ |
| | **Plus** | 0.021 | 1 | $10^{-314}$ |
| | **Only KIF16B** | $10^{-183}$ | $10^{-314}$ | 1 |

**Table 7 | Statistical comparisons of simulated negative and positive velocities of all dual-motor cargoes.** Supplementary to Fig. 5c and Extended Data Fig. 5b

| Motor composition and test | p-values with Bonferroni correction (3 comparisons) | | | | |
|---|---|---|---|---|---|
| 8 DDB + 6 KIF16B **Mann-Whitney U test** | | **Only DDB** | **Reversal pause** | **Non-Reversal pause** | **Only KIF16B** |
| | **Only DDB** | 1 | $10^{-125}$ | $10^{-220}$ | 0 |
| | **Reversal pause** | $10^{-125}$ | 1 | $10^{-14}$ | $10^{-121}$ |
| | **Non-Reversal pause** | $10^{-220}$ | $10^{-14}$ | 1 | $10^{-208}$ |
| | **Only KIF16B** | 0 | $10^{-121}$ | $10^{-208}$ | 1 |
| 8 DDB + 14 KIF16B **Mann-Whitney-U Test** | | **Only DDB** | **Reversal pause** | **Non-Reversal pause** | **Only KIF16B** |
| | **Only DDB** | 1 | $10^{-222}$ | $10^{-294}$ | 0 |
| | **Reversal pause** | $10^{-222}$ | 1 | $10^{-25}$ | $10^{-240}$ |
| | **Non-Reversal pause** | $10^{-294}$ | $10^{-25}$ | 1 | 0 |
| | **Only KIF16B** | 0 | $10^{-240}$ | 0 | 1 |
| 8 DDB + 20 KIF16B **Mann-Whitney-U Test** | | **Only DDB** | **Reversal pause** | **Non-Reversal pause** | **Only KIF16B** |
| | **Only DDB** | 1 | $10^{-236}$ | $10^{-307}$ | 0 |
| | **Reversal pause** | $10^{-236}$ | 1 | $10^{-24}$ | $10^{-248}$ |
| | **Non-Reversal pause** | $10^{-307}$ | $10^{-24}$ | 1 | 0 |
| | **Only KIF16B** | 0 | $10^{-248}$ | 0 | 1 |
| 8 DDB + 25 KIF16B **Mann-Whitney-U Test** | | **Only DDB** | **Reversal pause** | **Non-Reversal pause** | **Only KIF16B** |
| | **Only DDB** | 1 | $10^{-185}$ | $10^{-277}$ | 0 |
| | **Reversal pause** | $10^{-185}$ | 1 | $10^{-22}$ | $10^{-191}$ |
| | **Non-Reversal pause** | $10^{-277}$ | $10^{-22}$ | 1 | $10^{-291}$ |
| | **Only KIF16B** | 0 | $10^{-191}$ | $10^{-291}$ | 1 |

**Table 8 | Statistical comparisons of simulated reversal and non-reversal pause duration of reversal tracks for all dual-motor cargoes.** Supplementary to Extended Data Fig. 5c

| Motor composition and test | p-values | | |
|---|---|---|---|
| 8 DDB +<br>6 KIF16B<br>**Mann-Whitney-U Test** | | **Reversal pause** | **Non-Reversal pause** |
| | **Reversal pause** | 1 | 0.0002 |
| | **Non-Reversal pause** | 0.0002 | 1 |
| 8 DDB +<br>14 KIF16B<br>**Mann-Whitney-U Test** | | **Reversal pause** | **Non-Reversal pause** |
| | **Reversal pause** | 1 | $10^{-10}$ |
| | **Non-Reversal pause** | $10^{-10}$ | 1 |
| 8 DDB +<br>20 KIF16B<br>**Mann-Whitney-U Test** | | **Reversal pause** | **Non-Reversal pause** |
| | **Reversal pause** | 1 | $10^{-11}$ |
| | **Non-Reversal pause** | $10^{-11}$ | 1 |
| 8 DDB +<br>25 KIF16B<br>**Mann-Whitney-U Test** | | **Reversal pause** | **Non-Reversal pause** |
| | **Reversal pause** | 1 | $10^{-11}$ |
| | **Non-Reversal pause** | $10^{-11}$ | 1 |

**Table 9 | Statistical comparisons of reversal and non-reversal pause duration of reversal tracks for all simulated dual-motor cargoes without inactive motors.** Supplementary to Extended Data Fig. 6b

| Condition | Motor composition | Direction | Velocity (µm/s) (median ± IQR) | Number of cargoes |
|---|---|---|---|---|
| Conventional attachment rate | 8 DDB | Minus | -1.12 ± 0.46 | 142 |
| | 8 DDB + 14 KIF16B | Minus | -1.11 ± 0.72 | 73 |
| | | Reversal-Minus | -1.11 ± 0.73 | 228 |
| | | Reversal-Plus | 0.61 ± 0.25 | 228 |
| | | Plus | 0.62 ± 0.23 | 78 |
| | 14 KIF16B | Plus | 0.47 ± 0.14 | 148 |
| High attachment rate | 8 DDB | Minus | -1.07 ± 0.38 | 144 |
| | 8 DDB + 14 KIF16B | Minus | -0.37 ± 0.28 | 40 |
| | | Reversal-Minus | -0.39 ± 0.33 | 21 |
| | | Reversal-Plus | 0.35 ± 0.13 | 21 |
| | | Plus | 0.41 ± 0.19 | 97 |
| | 14 KIF16B | Plus | 0.41 ± 0.11 | 150 |
| Low detachment rate | 8 DDB | Minus | -1.19 ± 0.66 | 140 |
| | 8 DDB + 14 KIF16B | Minus | -0.86 ± 0.57 | 69 |
| | | Reversal-Minus | -0.99 ± 0.70 | 17 |
| | | Reversal-Plus | 0.62 ± 0.25 | 17 |
| | | Plus | 0.75 ± 0.28 | 55 |
| | 14 KIF16B | Plus | 0.58 ± 0.12 | 150 |
| High motor numbers | 128 DDB | Minus | -0.89 ± 0.19 | 150 |
| | 128 DDB + 230 KIF16B | Minus | -0.42 ± 0.30 | 60 |
| | | Reversal-Minus | -0.51 ± 0.26 | 32 |
| | | Reversal-Plus | 0.40 ± 0.17 | 32 |
| | | Plus | 0.42 ± 0.23 | 21 |
| | 230 KIF16B | Plus | 0.11 ± 0.01 | 150 |

**Table 10 | Median ± IQR of mean instantaneous velocities.** Values are given for minus-end directed runs (blue background) and plus-end directed runs (green background) of simulated single- and dual-motor cargoes with conventional parameter set, high attachment rates, low detachment rates and higher number of motors. Supplementary to Extended Data Fig. 7b

| Motor composition and test | p-values with Bonferroni correction (3 comparisons) - Mean velocities of negative runs- | | | |
|---|---|---|---|---|
| 8 DDB + 14 KIF16B High attachment rate **Mann-Whitney U test** | | **Only DDB** | **Minus** | **Reversal-Minus** |
| | **Only DDB** | 1 | $10^{-50}$ | $10^{-17}$ |
| | **Minus** | $10^{-50}$ | 1 | 1 |
| | **Reversal-Minus** | $10^{-17}$ | 1 | 1 |
| 8 DDB + 14 KIF16B Low detachment rate **Welch's test** | | **Only DDB** | **Minus** | **Reversal-Minus** |
| | **Only DDB** | 1 | $10^{-24}$ | 1 |
| | **Minus** | $10^{-24}$ | 1 | 0.019 |
| | **Reversal-Minus** | 1 | 0.019 | 1 |
| 128 DDB + 230 KIF16B **Mann-Whitney U test** | | **Only DDB** | **Minus** | **Reversal-Minus** |
| | **Only DDB** | 1 | $10^{-38}$ | $10^{-15}$ |
| | **Minus** | $10^{-38}$ | 1 | 0.779 |
| | **Reversal-Minus** | $10^{-15}$ | 0.779 | 1 |

| Motor composition and test | p-values with Bonferroni correction (3 comparisons) - Mean velocities of positive runs- | | | |
|---|---|---|---|---|
| 8 DDB + 14 KIF16B High attachment rate **Mann-Whitney U test** | | **Reversal-Plus** | **Plus** | **Only KIF16B** |
| | **Reversal-Plus** | 1 | 0.429 | 0.6317 |
| | **Plus** | 0.429 | 1 | 0.3397 |
| | **Only KIF16B** | 0.6317 | 0.3397 | 1 |
| 8 DDB + 14 KIF16B Low detachment rate **Mann-Whitney U test** | | **Reversal-Plus** | **Plus** | **Only KIF16B** |
| | **Reversal-Plus** | 1 | 0.016 | 1 |
| | **Plus** | 0.016 | 1 | $10^{-7}$ |
| | **Only KIF16B** | 1 | $10^{-7}$ | 1 |
| 128 DDB + 230 KIF16B **Welch's test** | | **Reversal-Plus** | **Plus** | **Only KIF16B** |
| | **Reversal-Plus** | 1 | 1 | $10^{-16}$ |
| | **Plus** | 1 | 1 | $10^{-13}$ |
| | **Only KIF16B** | $10^{-16}$ | $10^{-13}$ | 1 |

**Table 11 | Statistical comparisons of negative and positive velocities of all simulated dual-motor cargoes with high attachment rate, low detachment rate and higher number of motors.** Supplementary to Extended Data Fig. 7b

### Supplementary Videos

**Video 1 | Processive motility of DDB-eGFP**. Representative movie of DDB-eGFP complexes moving on a single microtubule (microtubule not shown). Minus end of the microtubule is located on the left. Movie plays 6xm faster than real time.

**Video 2 | Processive motility of KIF16B-eGFP**. Representative movie of KIF16B-eGFP motors moving on a single microtubule (microtubule not shown). Minus end of the microtubule is located on the left. Movie plays 6x faster than real time.

**Video 3 | Minus-end directed motility of DDB-vesicles**. Representative movie of DDB-vesicle (orange) moving on a single, polarity-marked microtubule (cyan). Minus end of the microtubule, represented by the brighter fluorescence signal, is located on the left. Movie plays 6x faster than real time.

**Video 4 | Plus-end directed motility of KIF16B-vesicles**. Representative movie of KIF16B-vesicle (orange) moving on a single, polarity-marked microtubule (cyan). Minus end of the microtubule, represented by the brighter fluorescence signal, is located on the left. Movie plays 6x faster than real time.

**Video 5 | Minus-end, plus-end and reversing motility of DDB-KIF16B-vesicles**. Representative movie of DDB-KIF16B-vesicle (orange) moving on a single, polarity-marked microtubule (cyan). Minus end of the microtubule, represented by the brighter fluorescence signal, is located on the left. Movie plays 6x faster than real time.

**Video 6 | DDB-KIF16B vesicle undergoing elongation during the paused states**. Example movie of DDB-KIF16B-vesicle (orange) moving on a single, polarity-marked microtubule (cyan) and undergoing elongation during the paused states. Minus end of the microtubule, represented by the brighter fluorescence signal, is located on the left. Movie plays 6x faster than real time.

### Materials and Methods

#### Reagents

Plasmids containing the gene for DIC2-SNAPf and p62-Halo were a kind gift from Prof. Samara Reck-Peterson, University of California San Diego[15]. Full length *H. sapiens* KIF16B gene in pFastBac vector was a kind gift from Prof. Marino Zerial, MPI-CBG, Dresden[16]. cDNA encoding the gene for *M. musculus* BICD2 was obtained from genomics-online (ABIN3826068). 18:1 (Δ9-Cis) DOPC (850375P), 18:1 (Δ9-Cis) DOPE (850725P), 18:1 PI3P (850150P), 18:1 DGS-NTA(Ni) (790404C) were obtained from Avanti-Polar while 18:1 DOPE-Atto647N (45-2247) was obtained from Millipore Sigma. Antibodies against dynein heavy chain (sc-514579), dynactin p62 (sc-55604), dynactin p150 (sc-135890), dynactin p50 (sc-393389) and dynactin Arp1 (sc-390632) were obtained from Santa Cruz, Texas; antibody against dynein intermediate chain (D5167) was obtained from Sigma-Aldrich; HRP-conjugated secondary antibody against mouse IgG (ab97023) was obtained from Abcam.

#### Molecular Cloning and baculovirus production

All plasmids were constructed by PCR and conventional restriction endonuclease and DNA ligation methods. Vector backbones with 3C protease cleavage sites and indicated fusion tags were provided by the protein expression, purification and chromatography (PEPC) facility at the Max Planck Institute, Cell Biology and Genetics, Dresden, Germany[17]. cDNA encoding DIC2-SNAPf and p62 were introduced into a vector downstream of the human cytomegalovirus (CMV) immediate-early promoter and enhancer with C-terminal affinity tags- MBP for DIC2-SNAP and TwinStrep for p62. A gBlock sequence (Integrated DNA Technologies) coding for 8xHistidine tag and streptavidin binding protein (SBP) in tandem was fused to the ORF corresponding to either BICD2N594 or BICD2N594-eGFP at the 3' end using High Fidelity assembly reaction (New England Biolabs) and introduced into an *E. coli* expression vector with an N-terminal MBP affinity tag. The gene coding for KIF16B was subcloned into an insect expression vector with and without a C-terminal e-GFP tag and an MBP affinity tag. All constructs contained a PreScission 3C protease site between the gene-of-interest and the affinity tags. All plasmids were verified by DNA sequencing.

Recombinant baculovirus for DIC2-SNAPf, p62 and KIF16B (with and without eGFP) were produced in Sf9 cells using the FlexiBAC system[17]. Briefly, plasmid vectors with the gene of interest were co-transfected with a replication-defective bacmid DNA into Sf9 cells. Homologous recombination between flanking sequences in both piece of DNA introduces the gene of interest into the viral genome and rescues viral replication and subsequent amplification. While recombinant viruses of P2 or P3 were used for DIC2-SNAPf and p62 (depending on the titre), P2 viruses were used for KIF16B. Viruses were either stored at 4 °C for one month or were supplemented with 10% (w/v) sucrose (in PBS; phosphate buffered saline, pH 7.3) and stored at – 80 °C for no more than 6 months.

**Protein purification and concentration estimation**

Dynein and dynactin: We exploited the ability of recombinant baculovirus to infect mammalian cells [18] to introduce affinity-tagged DIC2-SNAPf or p62 into HEK293F cells as bait for the dynein and dynactin complex, respectively. Genes for bait proteins in tandem with an affinity tag (Maltose binding protein, MBP for IC2 and Twin-Strep for p62) were introduced into large-scale suspension cultures of HEK293 via BacMam [18]. These bait proteins are incorporated into their respective complexes and can be fished out from cell lysates using affinity chromatography[15]. Peak expression of either bait was determined by performing a time course study on a small-scale suspension culture of HEK293F cells. The expression level for both dynein and dynactin was measured by running equal amount of cell lysate, sampled at 24 hr intervals, on a 4-12% BisTris SDS-PAGE precast gel in MOPS buffer (Life Technologies), which were then blotted onto a PVDF membrane (BioRad) and probed with either a primary anti-dynein intermediate chain or anti-dynactin p62 antibody and secondary anti-mouse IgG-HRP. Sufficient expression of p62 required the addition of sodium butyrate to a final concentration of 5 mM 6 hr post infection. Sodium butyrate inhibits histone deacetylases which, otherwise, suppress protein expression from extrachromosomal DNA[19–21]. Peak expression of both DIC2-SNAPf and p62 was observed at 24 hr post infection when infected with 1:100 and 1:50 P2 virus:culture (v/v) respectively.

To purify dynein or dynactin, 2-3 L of 2 x $10^6$ cells/mL HEK293F cells were infected with the respective baculovirus and grown in suspension for 24 hr at 37 °C and harvested by centrifugation. Cell pellets were washed with ice-cold PBS, resuspended in equal volumes of dynein lysis buffer (30 mM HEPES pH 7.4, 100 mM KCl, 2 mM $MgCl_2$, 1 mM EGTA, 10% glycerol, 0.02% Triton X-100, 0.2 mM MgATP, 1 mM DTT, 2 mM PMSF and 1X protease inhibitor cocktail (cOmplete, Roche) and lysed with one passage through an ice-cooled Avestin Emulsiflex C-5 homogenizer at 3000-5000 psi. The lysate was clarified at 200,000 xg for 60 min at 4 °C and the supernatant was then incubated with 2 mL of beads, equilibrated in dynein lysis buffer, for 120 min at 4 °C on a tube rotator. We used amylose resin (E8021S NEB) for dynein and streptactin beads (28-9355-99, Cytvia) for dynactin. Protein-bound beads were then collected with gravity flow 20 mL chromatography column (Econo-Pac, Bio-Rad), washed with 25 mL of dynein lysis buffer without PMSF and 25 mL of dynein elution buffer (30 mM HEPES pH 7.4, 148 mM potassium acetate, 2 mM magnesium acetate, 1 mM

EGTA, 10% Glycerol, 0.02% Triton X-100, 0.2 mM MgATP, 1 mM DTT). Dynein complexes were released from the amylose resin by cleaving the MBP tag with 20 µg/mL of PreScission 3C-Protease (diluted in the dynein elution buffer, obtained from PEPC facility, MPI-CBG) for 60 min at 4°C while dynactin was eluted from the streptactin resin with 2 mL of 2.5 mM d-Desthiobiotin (prepared in dynein elution buffer). Complexes were concentrated to ~ 500 µL with a 100K MWCo AmiconUltra (UFC8100, Amicon) centrifugal concentrator, filtered through a 0.22 µm cellulose acetate spin filter (Costar, Utah) and passed through a Superose 6 increase 10/300 GL size-exclusion column in dynein gel filtration (Composition?) buffer using a liquid chromatography system (ÄKTA pure). Peak fractions (corresponding to 11-13 mL retention volume) were pooled and concentrated to ~ 250 µL, aliquoted in volumes of 5 µL, snap-frozen and stored in liquid nitrogen.

*BICD2N594:* *M. musculus* bicaudal 2 (BICD2) truncated to first 594 amino acids was expressed in *E. coli* pRARE cells transformed with plasmid DNA carrying the gene in 750 mL Luria-Bertani medium containing 50 µg/mL kanamycin. The cells were grown at 37 °C with continuous shaking at 180 rpm till the optical density of 0.6 at 600 nm. Protein expression was induced by adding isopropylthio-β-galactoside (IPTG) to a final concentration of 0.5 mM and shaking the culture at 180 rpm at 18 °C for 18 hrs. The cells were then harvested at 7500 x g for 10 min at 18 °C and the pellets were resuspended in 35 mL BICD buffer (30 mM HEPES pH 7.4, 150 mM KCl, 2 mM MgCl2, 10% Glycerol, 0.02 % Triton X-100, 1 mM DTT) containing 1X protease inhibitor cocktail (cOmplete, Roche) and lysed by passaging it three times through an ice-cold Emulsiflex homogenizer at 5000-10000 psi. The lysate was supplemented with 10 µL Benzonase and spun at 186,000 x g at 4 °C for 45 min in a Type Ti-45 rotor (Beckman-Coulter) and the supernatant was passed through a 0.45 µm filter. The clarified lysate was incubated with pre-equilibrated amylose resin (one wash with water and two washes with BICD buffer) for 2 hr at 4 °C on a rotary mixer. The resin was collected in a fresh gravity flow column, washed with 50 mL BICD buffer, and eluted with 20 mM maltose (supplement in BICD buffer). Constructs were either directly processed for protease cleavage or subjected to nickel-sepharose affinity chromatography using the C-terminal 8xHis tag. Such tandem purification with affinity tags at the N and C termini allow the exclusion of truncated proteins from the final preparations. Eluted proteins were passed through a 1 mL HisTrap HP column (29051021, Cytiva), which was pre-equilibrated with 10 column volumes (CV) of BICD buffer containing 30 mM imidazole followed by a wash with 10 CV of 30 mM imidazole containing BICD buffer and elution with 300 mM imidazole containing BICD buffer. Eluted proteins were incubated with 5 µg/mL of PreScission 3C-protease at 4 °C overnight, concentrated using 30K MWCo spin filters and subjected to size-exclusion chromatography on a Superose 6 increase 10/300 column. Peak fractions (12-14 mL retention volume) were collected, concentrated, aliquoted, snap-frozen in liquid nitrogen and stored at -80 °C.

*KIF16B:* KIF16B was expressed and purified from Sf9 using baculovirus. Optimal expression conditions (infection ratio of 1:100 virus: culture, v/v, for 72 hr at 27 °C) were identified by performing a time-course experiment, similar to that described for DIC2-SNAPf and p62 above with one difference: protein expression in cell lysates was observed by Coomassie (LC6065, SafeStain) staining of SDS-PAGE gels. Fresh or snap-frozen cell pellets from 500 mL, 1 x $10^6$ cells/mL of Sf9 suspension culture infected with 5 mL of P2 virus carrying the gene for KIF16B were resuspended in 25 mL equilibration buffer (25 mM HEPES pH 7.2, 1 M NaCl, 5 mM $MgCl_2$, 5% Glycerol, 0.1 mM ATP, 0.25% 3-((3-cholamidopropyl) dimethylammonio)-1-propanesulfonate (CHAPS), 10 mM β-mercaptoethanol) supplemented with 1X protease inhibitor cocktail (cocktail III Merck calbiochem, 535140) and 1 µL Benzonase (1 mg/mL, MPI-CBG) to shear any DNA released during lysis. Cells were lysed by two passages through an ice-cold Avestin Emulsiflex C-5 homogenizer at 3000-5000 psi and spun at 186,000 x g for 60 min at 4 °C. The supernatant was filtered through a 0.45 µm syringe filter and incubated with 3 mL of pre-equilibrated amylose resin (one wash with distilled water and three washes with ice-cold lysis buffer, bead volume = 1.5 mL) for 2 hr on a rotary mixer in the cold room. Beads were collected in an empty gravity flow column, washed twice with 10 mL equilibration buffer, and eluted with 3 mL of equilibration buffer supplemented with 20 mM maltose. Eluted proteins were diluted to 10 mL with equilibration buffer and incubated with 5 µg/mL of PreScission 3C-protease (His-tagged) on a rotary mix overnight in the cold room. The protease was removed by incubating the mixture of protein and protease with 1 mL of pre-equilibrated Ni-NTA resin (30210, Qiagen) for 60 min followed by passage through a fresh gravity-flow column. The flow-through was concentrated in a 30K MWCo spin filters to ~1 ml, and gel-filtered on a Superose 6 increase 10/300 column in equilibration buffer at 4 °C. Peak fractions were pooled, and concentrated using fresh 30K MWCo spin filters, aliquoted, snap-frozen in liquid nitrogen and stored at -80 °C.

SDS-PAGE was used to determine the purity of isolated proteins and western blotting was used to confirm the identity of dynein heavy chain and dynein intermediate chain in dynein preparations and dynactin p150, dynactin p62, dynactin p50 and dynactin arp1 in dynactin preparations. Protein estimation was performed by running serial dilutions of a standard 6xHis-eGFP construct (MPI-CBG) alongside appropriate dilutions of the proteins on an 4-12% BisTris SDS-PAGE precast gel in MOPS or MES buffer (LifeTechnologies), stained with Coomassie for 60 min and de-stained in distilled water. Stained gels were imaged in an imaging station (c300, Azure Biosystems). The integrated intensity of the protein bands of interested was quantified using the 'Gels' tool in Fiji[22]. Individual lanes were defined using the rectangle region-of-interest tool, intensity profiles of all lanes were plotted, and integrated intensities were recorded by selecting the area under each peak. Linear fitting of the integrated intensity vs concentration of 6xHis-eGFP provided a calibration curve which was then used to estimate the concentration of desired proteins. The molar concentrations of DHC and p150 were defined as the concentration of dynein and dynactin respectively.

Tubulin: Porcine brain tubulin was purified by two rounds of polymerization and depolymerization in high molarity PIPES[23]. Rhodamine labelled tubulin was prepared using the labelling kit from Invitrogen following manufacturer's instructions.

**Preparation of vesicles**

All phospholipids were acquired as lyophilized powders and resuspended in chloroform except for PI3P where a 200:100:3 (v/v/v) mixture of chloroform:methanol:water was used a solvent. A thin film of phospholipids mixed in the molar ratio mentioned in supplementary table 1 was deposited on the inner walls of a clean glass vial under a light and steady stream of nitrogen followed by drying in a vacuum for 4 hr. Multilamellar vesicles (MLVs) were prepared by hydrating the lipid film in dynein buffer B (30 mM HEPES pH 7.4, 50 mM potassium acetate and 2 mM magnesium acetate) supplemented with 5% (w/v) sucrose to final lipid concentration of 1 mg/mL (1.28 mM) with vigorous shaking on a vortex and stored at -20 °C. MLVs with a total lipid mass of 50 µg were freeze-thawed five times followed by 10 cycles (21 passages) through an extruder (Avanti Polar) containing a phosphocellulose membrane with a pore size of 100 nm. Vesicles were stored on ice and used with 24-48 hr of preparation. Size distribution of vesicles (**Extended Data Fig. 1**) were measured in a dynamic light scattering instrument (Zetasizer, Malvern). The theoretical molar concentration of PI(3)P in the final vesicle solution was used as the molar concentration of vesicles.

**Preparation of polarity-marked microtubules**

Polarity-marked microtubules were prepared by preferably extending the plus-ends of short GMPCPP seeds in the presence of N-ethlymalemide modified tubulin (NEM-tubulin). Briefly, short bright seeds of 1:3 rhodamine labelled tubulin (final conc. 20 µM) were polymerized in the presence of 1 mM GMPCPP (NU-402, Jena Bioscience) in BRB80 for 15 min at room temperature. An extension mix comprising of 15 µM 1:9 rhodamine labelled tubulin, 6 µM fresh NEM-tubulin (40 µM unlabelled tubulin incubated with 1 mM NEM on ice for 10 min and excess NEM quenched with 10 mM DTT for 10 min), 2 mM GTP and 2 mM $MgCl_2$ in BRB80 was assembled on ice, warmed at 37 °C for 1 min and incubated with 1/20$^{th}$ volume of bright seeds at 37 °C for 20 min. Microtubules were stabilized with 10 µM taxol (Paclitaxel, Sigma) and harvested by spinning at 17,000 x g for 15 min. Polarity-marked microtubules were used within 72 hr.

#### Motility experiments

Water-tight flow channels were prepared from silanized glass coverslips (22 x 22 mm and 18 x 18 mm; cleaned in piranha solution and treated with 0.05% dichlorodimethylsilane) by placing 1.5 mm parafilm strips on a 22 x 22 mm coverslip (~ 3 mm) apart, covering with an 18 x 18 mm coverslip, and melting the parafilm at 55 °C heat block. All solutions were prepared in dynein buffer B. The channels were sequentially perfused with solutions containing TetraSpeck microspheres (diameter 0.1 µm, Invitrogen) for drift and color correction, monoclonal anti-β-tubulin antibodies to immobilize microtubules and 1% Pluronic F127 to block the surface. After a 60 min incubation, channels were washed twice with dynein buffer B and once with dynein buffer B containing 10 µM taxol and incubated with polarity-marked microtubules for 5 min.

For single-molecule DDB motility, 50 nM dynein and 100 nM dynactin were first incubated on ice for 5 min (= 50 nM DD). 38 nM DD was then mixed with 75 nM BICD2N594-eGFP in dilution buffer (dynein buffer B supplemented with 20 mM glucose, 0.1 mg/mL casein, 2.5 mM MgATP and 1 mM DTT) and incubated for 5 min on ice (= 38 nM DDB). DDB complexes were diluted 4-fold with the imaging buffer (dynein buffer B supplemented with 20 mM glucose, 0.1 mg/mL casein, 2.5 mM MgATP, 1 mM DTT, 100 µg/mL glucose oxidase and 20 µg/mL catalase) and perfused into flow channels. Single molecule motility of KIF1B-eGFP was observed by serially diluting the stock solution of KIF16B-eGFP to a final concentration of 1 nM in imaging buffer and perfusing into the flow channel.

Dual-motor vesicle assays were performed by incubating 120 nM vesicles with 840 nM BICD2N594 and different concentrations of KIF6B (mentioned in the text) or 25 nM KIF16B-eGFP (to detect the presence of KIF16B on minus-end directed and reversing vesicles) for 2 min on ice. 24 nM of these vesicles were then incubated with 38 nM DD for 5 min on ice, diluted 4-fold in imaging buffer and perfused into the flow channels. Single-motor vesicle assays were performed by replacing either KIF16B (for DDB-vesicles) or DDB (for KIF16B-vesicles) from the above mixture with dilution buffer.

#### Data acquisition

TIRF Microscopy was performed on a Nikon Eclipse Ti2 microscope equipped with a perfect focus system and a 100X, 1.49 NA oil, apochromat TIRF objective and 1.5X optovar. Samples were illuminated with either 488 nm, 561 nm or 647 nm lasers (100 mW each) placed in a visitron laserbox and channeled through an iLas2 ring TIRF module operated in a single angle mode (point TIRF). Images from different fluorescent channels were acquired with separate EMCCD cameras (one iXon Life EMCCD each for 488 nm and 561 nm channels and iXon Ultra EMCCD for 647 nm channel) each containing 1024 x1024 pixel sensor and controlled with VisiView. The size of each pixel was 87 x 87 nm. Images were either acquired in streaming mode with 100 ms exposure (10.0 frames per second) or every 300 ms in time lapse mode with 100 ms exposure (3.3 frames per second).

**Data processing and analysis**

Tracking: Single molecules of DDB-eGFP and KIF16B-eGFP, and vesicles were tracked with Fluorescence Image Evaluation Software for Tracking and Analysis (FIESTA)[24] which automates Gaussian fitting of fluorescence signals to extract X-Y position coordinates. All tracks were manually curated to exclude erroneous tracks from further analysis. Tracks from vesicles at microtubule junctions and from vesicles which remained stationary throughout were discarded. Any tracks from vesicles colliding into each other and slowing down as a result were also excluded. Tracks were corrected for drift and chromatic aberration (wherever applicable) by using TetraSpeck microspheres (Invitrogen) as fiduciary markers. The X-Y position trace of individual tracks was used as a path to calculate displacement along path. This was necessary to avoid false directional reversals when simple a distance to origin is calculated for vesicles moving on curved microtubules. The orientation of tracks was adjusted such that motion towards the plus-end of the microtubule reflected as positive displacement.

Segmentation: Individual tracks (displacement along path vs time) were split into segments of "runs" and "pauses" (which also includes diffusion). Mean-square displacement method[25] was performed to identify these states. The data was first smoothened using a moving window average of 0.6 seconds. Mean-square displacements were calculated as a function of the lag time $\tau$ within a moving window of 6 seconds. The mean-square displacement $\langle x^2 \rangle$ as function of lag-time $\tau$ was fit with $\langle x^2(\tau) \rangle \propto \tau^{\alpha}$ to obtain the value of the scaling exponent α. Only $\tau < 3$ s was considered since less data points are available for longer lag times. The value of α determines if the motion within a window is either diffusive (α≈1), sub-diffusive (α <1), super-diffusive α >1) or ballistic (α≈2). The motion within a window is considered active (and therefore a run) if α > 1.7 [25]. If α < 1.7, the central data point of the window was assigned a paused state. To improve the segmentation, segments with only a single data point were merged with the next segment. Runs were classified as either negative or positive based on the slope of a cubic spline interpolation: negative slopes indicated negative runs and positive slopes indicated positive runs. Additional refinements were made for accurate detection of pauses. Runs with net displacement smaller than 300 nm were classified as pauses to reliably detect short pauses, which were otherwise misclassified due to data smoothening. Pauses that were erroneously detected at the beginning and the end of the track due to lower accuracies of the mean-square and cubic spline interpolations in these regimes were discarded. Only pauses that followed a run were considered for further analysis as there was no ambiguity about the activity of the motors.

*Velocity, spatial pause frequency, pause duration:* Instantaneous velocity was defined as the ratio of change in displacement and time between two consecutive frames. Velocity of positive and negative runs was defined as the arithmetic mean of instantaneous velocities of individual runs.

Spatial pause frequencies, $x_f$, of individual tracks were calculated as $x_f^i = \frac{x_p^i}{x_d^i}$, where $x_p^i$ is the number of pauses and $x_d^i$ the absolute distance traveled (absolute value for minus runs) of an individual track $i$. The spatial pause frequency of individual tracks was weighted by the proportion of total distance travelled by vesicles of a given type (minus, plus or reversal tracks), $w_i = \frac{x_d^i}{\sum_{i=0}^{N} x_d^i}$, to calculate the weighted mean spatial pause frequency (or simply pause frequency, $\overline{x_f}$), weighted standard deviation ($\sigma_f$) and weighted standard error of mean ($\sigma_{\bar{x}}$).

$$\overline{x_f} = \sum_{i=0}^{N} w_i . x_f^i \ , \ \sigma_f = \sqrt{\frac{\sum_{i=0}^{N} w_i \left(x_f^i - \overline{x_f}\right)^2}{\left(\frac{N-1}{N}\right)}} \ , \ \sigma_{\bar{x}} = \frac{\sigma_f}{\sqrt{N-1}}$$

$N$ is the total number of tracks. Pause duration was simply defined as the time interval between the beginning and end of a pause segment.

*Statistics:* Independent t-tests and Mann-Whitney-U tests were performed using the python module *scipy.stats*[26]. Bonferroni corrections of p-values were performed manually by multiplying the p-values given by the tests by the number of comparisons made. The two sample weighted KS-test was manually implemented based on the source code of two sample KS-test from *scipy.stats (*function *"ks_2samp"* in file *scipy/stats/stats.py*). Instead of the unweighted empirical cumulative distribution function

$$F(x_i) = \sum_{j=1}^{i} 1/N = i/N$$

the weighted empirical cumulative distribution function

$$F^w(x_i) = \sum_{j=1}^{i} w_j$$

with weights $w_j$ was used.

*Data representation:* The Beeswarm plot was generated using the python package *seaborn*[27]. All other plots were generated using the python package *matplotlib*[28] and formatted in Inkscape. Kymographs and timelapse images were created with Fiji[22].

**Mathematical modelling**

The mathematical model used in this work is based on previously published models[8,29,30]and has been adapted to the experimental geometry.

In the model, we assign a fixed number of motors to each cargo. It is expected that in the experiment, the number of motors as well as the composition of KIF16B and DDB motors slightly fluctuates between cargoes for a given concentration. To mimic these fluctuations, the number of motors (including active, inactive and diffusive DDB and KIF16B) are thrown from a Gaussian distribution with the mean being the given mean number of motors. The standard deviation of the Gaussian is a function of the mean:

$$\sigma(\mu) = 1.0683 \sqrt{\mu}$$

This relation was found in an extra simulation, where varying numbers of motors were randomly distributed over a fixed number of cargoes. Having the number of motors, the actual number of KIF16B and DDB is randomly generated with probabilities given by the mean numbers of DDB and KIF16B. Thus, beside the number of motors also the ratio of KIF16B to DDB can slightly vary between cargoes for the same given mean numbers of KIF16B and DDB.

On the cargo, motors are divided into an attachment area and a reservoir. Motors in the attachment area can attach the MT, while motors in the reservoir are too far away to interact with the MT. In our model, we only take the motors in the attachment area into account but let them exchange with the motors in the reservoir. This means, each time when a motor attaches, it is randomly chosen whether the motor is active, inactive or diffusive (in case of DDB, see later for the definitions of active, inactive and diffusive motors). Furthermore, the motor is assigned an individual maximal stepping rate upon attachment (see later for definition of motor stepping rates). However, the ratio between DDB and KIF16B, which was assigned previously to this particular cargo, is fixed.

The microtubule is modeled as one-dimensional lattice (one-dimensional coordinate system) with seven parallel lanes, which are representing the protofilaments accessible to the motors, i.e. the upper half of the microtubule. KIF16B and DDB motors, which are bound to the cargo, can bind to the microtubule, step on it and detach from the microtubule. Both KIF16B and DDB motors bind to the microtubule with constant attachment rates $k_{a,KIF16B}$ and $k_{a,DDB}$, respectively (see **Extended Data Tables 3 and 4** for parameter values and references). When attaching to the microtubule, a Gaussian distribution, peaked around the central protofilament, is used to randomly chose the lane (protofilament) the motor interacts with. The Gaussian distribution is cut at $\pm 3\sigma$, where standard deviation sigma is one and the mean is zero (**Extended Data Figs. 8)**. Once attached to a lane, the motors stay on this lane until they detach again.

When bound to the microtubule, motors exert a force on the cargo proportional to the motor deflection:

$$\Delta x^i(t) = x^i_{mt}(t) - X_V(t)$$

where $x^i_{mt}(t)$ is the motor head position on the microtubule and $X_V(t)$ the cargo (vesicle) position in the one-dimensional coordinate system parallel to the protofilament axis. The deflection does not depend on the selected lane.

The motors are modeled as Hookean springs with non-zero rest length $L_{KIF16B}$ and $L_{DDB}$, respectively. This means the forces, the motors exert on the cargo are given by

$$F^i(t) = \begin{cases} \kappa_{KIF16B}(\Delta x^i(t) - L_{KIF16B}), \text{ if } (\Delta x^i(t) > L_{KIF16B} \\ \kappa_{KIF16B}(\Delta x^i(t) + L_{KIF16B}), \text{ if } (\Delta x^i(t) < -L_{KIF16B} \\ 0, else \end{cases}$$

for KIF16B and by

$$F^i(t) = \begin{cases} \kappa_{DDB}(\Delta x^i(t) - L_{DDB}), \text{ if } (\Delta x^i(t) > L_{DDB} \\ \kappa_{DDB}(\Delta x^i(t) + L_{DDB}), \text{ if } (\Delta x^i(t) < -L_{DDB} \\ 0, else \end{cases}$$

for DDB. $\kappa_{KIF16B}$ and $\kappa_{DDB}$ are thereby the KIF16B and DDB specific motor stiffnesses, respectively.

For individual maximal stepping rate under zero load, experimentally measured single molecule KIF16B and DDB instantaneous velocities were used (**Fig. 1C** of the main text). For stepping under load, the stepping rate depends on the force regime. If the motor experiences a resisting force ($F^i(t)$>0 for KIF16B and $F^i(t)$<0 for DDB) smaller than the stall force ($F_{s,KIF16B} > F^i(t)$>0 for KIF16B and $-F_{s,DDB} < F^i(t)$<0), the stepping rate depends on the force and the ATP concentration as suggested by Schnitzer et al. 2000[31]:

$$s([ATP], F^i) = \frac{V_{max} \times [ATP]}{[ATP] + K_M} = \frac{k_{cat}(F^i) \times [ATP]}{[ATP] + k_{cat}(F^i)/k_b(F^i)}$$

with $k_{cat}(F^i)$ and $k_b(F^i)$ being Boltzmann-like distributed:

$$k_m(F^i) = \frac{k_m^0}{p_m + q_m e^{F^i\delta/k_BT}} \quad with\, m \in \{cat, b\}$$

While $k_{cat}^0 = v_f/d$ is determined by the motor forward velocity $v_f$ and step size $d$, the parameters $k_b^0$, $p_b + q_b = 1$ and $p_{cat} + q_{cat} = 1$ are taken from Schnitzer et al. 2000[31]. $\delta$ is determined by setting the stepping rate at the stall force equal $0.1\, s^{-1}$, i.e., $s(ATP, F_s^i) = 0.1\, s^{-1}$.

Under assisting forces ($F^i(t)$<0 for KIF16B and $F^i(t)$>0 for DDB), the stepping rate is equal to the force-dependent stepping rate at zero load: $s([ATP], F^i) = s([ATP], F^i = 0)$. If the motor experiences resisting forces beyond the stall force ($F_{s,KIF16B} < F^i(t)$ for KIF16B and $-F_{s,DDB} > F^i(t)$ for DDB), the motor steps backwards with a small but constant rate $s = v_b/d$.

While the formulas for the stepping rates are the same for KIF16B and DDB, the parameters such as forward velocity $v_f$ and stall force $F_s$ are different for KIF16B and DDB. Different stall forces and forward velocities lead to different force and ATP dependences for KIF16B and DDB. Even though the used force and ATP-dependent stepping was found for kinesin-1, the previously found force dependence of KIF16B and DDB are similar[32,33].

For stepping and attachment, steric motor hindrances (exclusion effects) are taken into account on the protofilaments. This means, a motor can neither attach nor step to an occupied spot on the same lane of the microtubule.

For KIF16B and DDB detachment, an exponentially-increasing detachment rate is used following previous work[4,7,11]:

$$k_d(F^i) = k_d^0\, e^{-|F^i|/F_d}$$

Different force-free detachment rates and detachment forces were used for KIF16B and DDB, respectively. The used force-free detachment rates $k_d^0$, were taken from experimentally measured run length and velocities of single KIF16B and DDB molecules, respectively.

Moreover, in single molecule experiments, it was found that about 20 % of KIF16B motors do not step at all. Therefore, the model also includes 20 % inactive KIF16B motors, which do not step when being attached to the microtubule but rather stay strongly bound. Thus, we assign them a lower force-free detachment rate than for the active motors. For DDB, it was found experimentally that about 10% do not step at all and about 10% diffuse along the MT. Thus 10 % of DDB motors are modeled as inactive motors with lower force-free detachment rate and 10 % of DDB motors are modeled to diffuse in the harmonic potential of the motor spring with the following rate

$$s_{\pm}(F^i) = s_0\, e^{\mp|F^i|\cdot d/2k_BT}$$

The diffusing motors hence always tend to step towards their equilibrium positions where they are not under tension.

The Gillespie stochastic simulation algorithm[34] is used to advance the system in time. The motion of the cargo is modeled in the over-damped limit, which is in agreement with experimental conditions. This means, after each motor update the cargo diffuses in the harmonic potential of attached motor springs around its equilibrium position, i.e., the position where forces exerted on the cargo are equilibrated. For the cargo diffusion the Metropolis algorithm is used[35]. The simulation starts with no motor being attached to the microtubule. The measurement begins after a relaxation time of 4 s. The simulation is terminated either when no motor is attached, or after 80 s. Number of simulated cargoes are provided in respective figures. Model parameters were either obtained from experiments or from literature, if possible. Certain parameter values obtained from literature as well as some unknown parameter values were optimized in order to fit the simulation best to the experiment. We made sure that parameter values remain in the range of those given in the literature. All parameters for DDB and KIF16B used in this work are summarized in **Extended Data Tables 4** and **5** respectively.
